# Cholinergic relay from punishment- to reward-encoding dopamine neurons signals punishment withdrawal as reward in *Drosophila*

**DOI:** 10.1101/2020.06.22.165209

**Authors:** Li Yan McCurdy, Preeti Sareen, Pasha A. Davoudian, Michael N. Nitabach

**Affiliations:** Interdepartmental Neuroscience Program, Yale University, New Haven, CT, USA; Department of Cellular & Molecular Physiology, Yale University, New Haven, CT, USA; Department of Neuroscience, Yale University, New Haven, CT, USA; MD/PhD Program, Yale School of Medicine, Yale University, New Haven, CT, USA; Department of Genetics, Yale University, New Haven, CT, USA

**Keywords:** Learning and memory, *Drosophila*, dopamine, mushroom body microcircuits

## Abstract

Animals form and update learned associations between otherwise neutral cues and aversive outcomes to predict and avoid danger in changing environments. When a cue later occurs without punishment, this unexpected withdrawal of aversive outcome is encoded as reward, via activation of reward-encoding dopaminergic neurons. Using real-time *in vivo* functional imaging, optogenetics, behavioral analysis, and electron-microscopy, we identify the neural mechanism through which *Drosophila* reward-encoding dopaminergic neurons are activated when an olfactory cue is unexpectedly no longer paired with electric shock punishment. Reduced activation of punishment-encoding dopaminergic neurons relieves depression of synaptic inputs to cholinergic neurons, which in turn synaptically increase odor responses of reward-encoding dopaminergic neurons to decrease odor avoidance. These studies reveal for the first time how an indirect excitatory cholinergic synaptic relay from punishment- to reward-encoding dopaminergic neurons encodes the absence of a negative as a positive, revealing a general circuit motif for unlearning aversive memories that could be present in mammals.

## INTRODUCTION

Animals use associative learning to build internal models of their environments for guiding adaptive decision making. They also flexibly update learned associations when new or conflicting information arises. Reversal learning paradigms probe cognitive flexibility in the context of changing stimulus-outcome contingencies (Costa et al., 2015; Izquierdo et al., 2017). Reversal learning comprises two components: acquisition and reversal. During acquisition, the “conditioned stimulus +” (CS+) cue is paired with a reinforcer (e.g., electric shock), while the distinct CS-cue is presented without the reinforcer, to form a specific association between CS+ and reinforcer. Later, during reversal, the stimulus-outcome contingencies are reversed: the reinforcer is omitted during CS+ presentation, and is instead delivered during CS-. Reversal learning requires cognitive flexibility in order for internal representation of each cue to be first assigned and later updated as contingencies change.

Extinction is the unlearning of the association between CS+ and reinforcer, such as occurs during reversal. Mammalian studies implicate dopaminergic neurons (DANs) in both the acquisition and extinction of aversive memories (Holtzman-Assif et al., 2010; Raczka et al., 2011; Badrinarayan et al., 2012; Luo et al., 2018; Salinas-Hernandez et al., 2018). Specifically, manipulation of dopamine signaling in the nucleus accumbens, prefrontal cortex or striatum interferes with reversal learning (Cools et al., 2007; Delamater, 2007; De Steno and Schmauss, 2009; Clarke et al., 2011; Hamilton and Brigman, 2015; Izquierdo et al., 2017). These studies suggest a key role for DANs in encoding the unexpected omission of aversive outcomes that drives extinction of cue-outcome associations. However, how withdrawal of reinforcer leads to changes in DAN activity remains unknown, as do the specific distinct roles possibly played by different subsets of DANs.

Here we use *Drosophila* flies as a model system to address these questions. *Drosophila* are capable of both forming and reversing cue-outcome associations, such as odor-shock associations, and have well-characterized neural circuitry and genetic tools for precise manipulation and visualization of neural activity at cellular resolution. The main brain structure underlying learning and memory in the fly is the mushroom body (MB) (Tully et al., 1990; Heisenberg, 2003; Schwaerzel et al., 2003; McGuire et al., 2005; Fiala, 2007; Keene and Waddell, 2007). Olfactory cues are represented by activity of sparse subpopulations of cholinergic Kenyon cell interneurons (KCs), which receive synaptic inputs from second-order olfactory projection neurons and innervate the neuropil of the MB (Aso et al., 2009; Aso et al., 2014a). MB output neurons (MBONs), comprising 21 subsets defined by the twenty anatomical MB compartments their dendrites innervate to receive KC synaptic inputs, project to downstream centers of the brain to promote approach or avoidance (Lin et al., 2007; Tanaka et al., 2008; Aso et al., 2014b; Owald et al., 2015). Approximately 130 modulatory DANs comprising 20 subsets distinguishable genetically and anatomically innervate the twenty compartments of the MB lobes (Tanaka et al., 2008; Mao and Davis, 2009; Aso et al., 2014a; Hige et al., 2015; Aso and Rubin, 2016). MBONs and DANs are named based on the MB lobe compartments they innervate, by reference to the α, β, and γ lobes and contiguous position along each lobe (Crittenden et al., 1998; Ito et al., 1998). DANs are anatomically classified as PPL1 or PAM DANs, which are viewed generally as encoding negative and positive reinforcement signals respectively (Schwaerzel et al., 2003; Riemensperger et al., 2005; Claridge-Chang et al., 2009; Mao and Davis, 2009; Liu et al., 2012; Das et al., 2014; Lin et al., 2014; Kirkhart and Scott, 2015). DANs secrete dopamine into the specific compartments they innervate, where they induce plasticity of KC-MBON synapses, thus modulating MBON odor responses and odor-evoked behavior (Sejourne et al., 2011; Pai et al., 2013; Placais et al., 2013; Aso et al., 2014b; Bouzaiane et al., 2015; Cohn et al., 2015; Hige et al., 2015; Owald et al., 2015; Perisse et al., 2016; Hattori et al., 2017; Berry et al., 2018; Handler et al., 2019; Zhang et al., 2019).

In *Drosophila*, as in mammals, reward-encoding DANs are thought to underlie the extinction of aversive memories, as silencing reward-encoding DANs using a broadly-expressed PAM genetic driver impairs extinction (Felsenberg et al., 2018). While much is known about learning-induced changes in odor-evoked neural activity in MBONs (Sejourne et al., 2011; Pai et al., 2013; Placais et al., 2013; Bouzaiane et al., 2015; Cohn et al., 2015; Hige et al., 2015; Owald et al., 2015; Perisse et al., 2016; Hattori et al., 2017; Berry et al., 2018; Handler et al., 2019; Zhang et al., 2019), few studies have assessed changes in DAN odor responses during acquisition (Riemensperger et al., 2005; Mao and Davis, 2009; Dylla et al., 2017), and none during reversal. DANs respond to various positive and negative reinforcers such as sugar, electric shock, and noxious heat, but it remains unknown in any model organism or learning paradigm how DAN signals are generated in response to withdrawal of negative reinforcement, nor how the absence of negative reinforcement is encoded as positive reinforcement.

Here we use a combination of functional imaging, optogenetics, and behavioral approaches in *Drosophila* to answer these key issues in the biology of reversal learning, specifically the neural mechanisms of extinction of CS+-shock association during reversal. We establish that PAM-β’2a DANs encode shock withdrawal during reversal as rewarding. This dopaminergic reward signal extinguishes CS+-shock association, and hence reduces CS+ avoidance, by depressing KC synapses onto avoidance-encoding MBON-γ5β’2a. We further demonstrate that approach-encoding MBON-γ2α’1 is an upstream element that activates PAM-β’2a when shock is withdrawn. Finally, we show that PPL1-γ2α’1 shock-responsive DANs relieve synaptic depression of KC-MBON-γ2α’1 synapses as a consequence of shock withdrawal to increase CS+ odor activation of MBON-γ2α’1. These studies reveal the underlying cellular and synaptic mechanisms through which DANs of opposite valence participate in an indirect relay from the γ2α’1 compartment to the β’2a compartment to encode withdrawal of a negative reinforcer as a positive reinforcer.

## RESULTS

### Reward encoding of shock withdrawal by PAM-β’2a DAN

To identify specific DAN subsets involved in reversal learning, we developed an experimental preparation for real-time recording of neural activity in genetically targeted neurons during aversive olfactory conditioning. We simultaneously expressed green GCaMP6f Ca^2+^ indicator and red Ca^2+^-insensitive tdTomato in neurons of interest for ratiometric visualization of Ca^2+^ changes reflecting neural activity. Each fly is head-fixed for simultaneous delivery of odors and electric shocks while recording neural activity (**Figure 1A**). During acquisition, flies receive alternating 5s pulses of the conditioned stimulus odor paired with electric shock (CS+) and a control odor (CS-), repeated for five trials (**Figure 1B**). During reversal, the contingencies are reversed for two trials: CS+ odor is presented without shock, and CS-odor is presented with shock.

**Figure 1:**
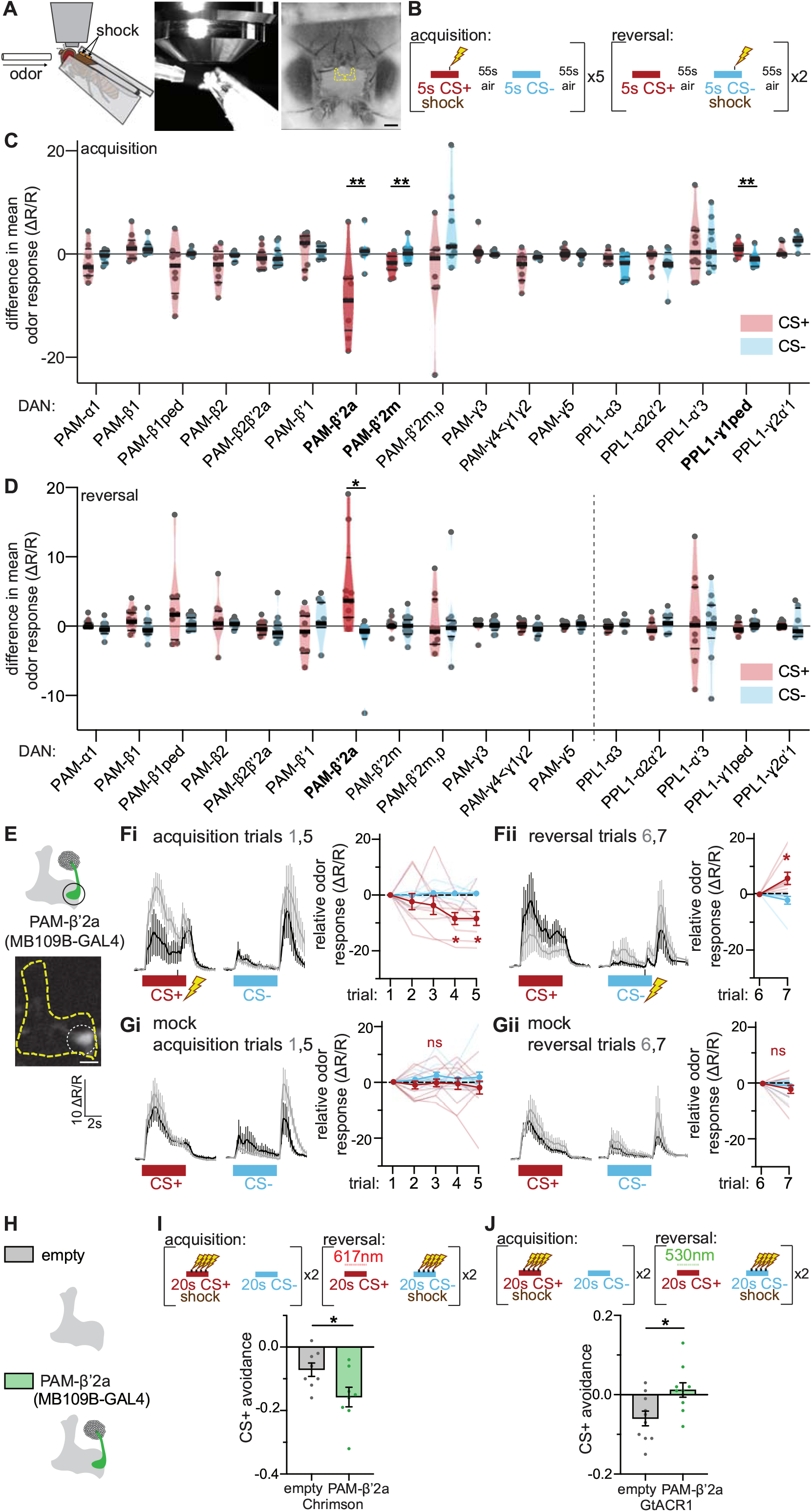
PAM-β’2a DANs encode reward of shock withdrawal during reversal learning. (**A**) Left: Setup for visualizing neural activity via Ca^2+^ imaging during aversive olfactory memory acquisition and reversal. Flies are head-fixed and cuticle dissected for ratiometric imaging of Ca^2+^-sensitive GCaMP6f and Ca^2+^-insensitive tdTomato. Middle: Photo of fly in imaging setup. Right: View of fly head and dissection window using a 5x objective; scale bar is 100μm. Location of MBs outlined in yellow. (**B**) Training paradigm for aversive memory acquisition and reversal. In five acquisition trials, 5s pulse of CS+ odor is paired with 100ms electric shock, and alternated with 5s pulse of CS-odor without shock; each odor pulse is separated by 55s clean air. Acquisition trials are followed by two reversal trials in which shock is omitted during CS+ odor presentation, and CS-odor is paired with shock. 4-methylcyclohexanol and 3-octanol were used as CS+ and CS-odors for the imaging screen respectively. (**C**) DAN imaging screen: acquisition trials. Relative difference from last to first acquisition trials in mean Ca^2+^ response of indicated PAM DANs to CS+ and CS-odors. Positive values indicate increase in odor response during acquisition. Each grey dots represents data from a single fly. Violin plots show range of data points, with median (thick black line) and quartiles (thin black lines). Change in PAM-β’2a, PAM-β’2m, and PPL1-γ1ped CS+ odor response relative to CS-odor responses are significantly different. *n* = 7-12 flies per genotype. Statistical comparisons by paired *t*-test or Wilcoxon matched-pairs signed rank test where appropriate, between CS+ and CS-odor response for each genotype. (**D**) DAN imaging screen: reversal trials. Same as in **C**, except for reversal trials. Change in PAM-β’2a CS+ odor response relative to CS-odor response is significantly different. (**E**) Schematic and sample fluorescence image of PAM-β’2a, the only DAN that exhibits a significant change in CS+ odor response after a single reversal trial. Only one hemisphere depicted for clarity. Sample image is at 20x; scale bar is 20μm. Scale bar of ΔR/R applies to all mean fluorescence traces in this figure. (**F**) Neural activity of PAM-β’2a in flies undergoing aversive memory acquisition (**i**) and reversal (**ii**) trials. Left: Mean fluorescence traces, the first acquisition or reversal trial in grey, the last acquisition or reversal trial in black. Right: Line graphs show relative difference of mean odor response from first acquisition or reversal trial for individual flies (thin lines) and group means (thick lines). Mean Ca^2+^ odor response is quantified as average activity during the first 4s from odor onset. Positive values indicate increase in odor response. PAM-β’2a CS+ odor response decreases during acquisition and increases during reversal. *n* = 9 flies. Statistical comparison for acquisition is by repeated-measures two-way ANOVA with Dunnett’s post-hoc test relative to the first acquisition trial. Statistical comparison for reversal is by paired *t*-test. These statistical comparisons are used in all imaging figures where a single genotype is shown. Since the variance in data for CS+ and CS-differed substantially, a Greenhouse-Geisser correction is applied. (**G**) Same as in (**F**), but from flies undergoing mock aversive memory acquisition and reversal, i.e., odors are presented as in (**F**), but no shocks are delivered. No change in PAM-β’2a odor response occurs during acquisition or reversal. *n* = 11 flies. Same statistical comparisons are used as in (**F**). (**H**) Schematic of PAM-β’2a and empty-splitGAL4 control driver line used in behavioral experiments. (**I**) Top: Reversal protocol for behavioral experiments. During acquisition, 20s of CS+ odor is paired with four electric shocks, followed by 20s of CS-odor without reinforcement, with 20s of clean air presented in between each odor; this is done twice. After acquisition, flies undergo two reversal trials, in which 20s of CS+ odor is presented without electric shock, and 20s of CS-odor is presented with electric shock. Red light illuminates the shock tube during CS+ presentation during reversal to activate neurons expressing Chrimson. Flies are then placed in quadrant arena and odor preference (CS+ vs CS-) is determined. CS+ odor avoidance was quantified as the fraction of flies in the CS-quadrants after being allowed to explore for two minutes; a value of 1 indicates that 100% of the flies avoided CS+ quadrants, a value of 0 indicates that flies chose CS+ and CS-quadrants equally. 3-octanol and 4-methylcyclohexanol were used as CS+ and CS-odors in separate groups of flies trained and tested contemporaneously and averaged to generate a reciprocally balanced preference score. Bottom: Control flies show slight preference for CS+ quadrants. Activating PAM-β’2a during CS+ reversal decreases CS+ avoidance relative to genetic controls, indicating enhanced extinction of aversive CS+ memory. *n* = 8 independent groups of flies per genotype. Statistical comparison is unpaired *t*-test. (**J**) Top: Same protocol as in (**I**), except using green light to silence neurons expressing GtACR1. Bottom: Silencing PAM-β’2a during CS+ reversal impairs extinction of aversive CS+ memory, reflected in a relative increase in CS+ avoidance. *n* = 10 independent groups of flies per genotype. Statistical comparison is by unpaired *t*-test. For all graphs, error bars are mean ± s.e.m, * *p*<0.05, ** *p*<0.01, *** *p*<0.001, **** *p*<0.0001.

Using this approach, we systematically characterized dynamic changes in CS+ and CS-odor response during acquisition and reversal of each DAN subset targeted individually using a library of intersectional “split-GAL4” driver lines (Jenett et al., 2012; Aso et al., 2014a). For this screen, we used 4-methyl-cyclohexanol as CS+ odor and 3-octanol as CS-odor. We identified two of seventeen DAN subsets that decrease CS+ odor response relative to CS-odor response during acquisition: PAM-β’2a and PAM-β’2m (**Figure 1C**). PPL1-γ1ped increases CS+ odor response during acquisition, as previously observed (Dylla et al., 2017). While it decreases during acquisition, PAM-β’2a DAN increases CS+ odor response during reversal (**Figure 1D**). No other DANs changes their odor responses during acquisition or reversal.

PAM-β’2a significantly decreases its CS+ odor response only after three acquisition trials, yet significantly increases its CS+ odor response after a single reversal trial (**Figure 1E**,**F**). These changes were replicated in a separate group of flies using reciprocal odor identities: 3-octanol and 4-methylcyclohexanol as the CS+ and CS-odors, respectively (**Figure S1B**). Importantly, no changes in odor response occur during mock acquisition or reversal, in which no shocks are delivered, indicating that changes in odor response are dependent on odor-shock pairing (**Figure 1G, S1C**).

Because PAM DANs are traditionally considered to encode reward (Liu et al., 2012; Das et al., 2014; Lin et al., 2014; Lewis et al., 2015), PAM-β’2a’s increase in CS+ odor response during reversal suggests that it could encode the rewarding value of shock withdrawal. To test this, we optogenetically interrogated the functional role of PAM-β’2a during a behavioral reversal learning paradigm (**Figure 1I, S1D**), in which groups of flies are trained with two acquisition trials and two reversal trials in a standard shock-odor delivery chamber and then tested for their odor preference in a four-quadrant two-odor choice chamber (Aso et al., 2014b; Klapoetke et al., 2014). Optogenetic activation of PAM-β’2a using Chrimson red-light-gated ion channel (Klapoetke et al., 2014) during CS+ presentation in reversal trials (when shock is omitted) decreases CS+ avoidance, relative to control “empty-splitGAL4” flies not expressing Chrimson during subsequent testing (**Figure 1I**). This indicates that PAM-β’2a activity during CS+ presentation in reversal trials is sufficient to enhance extinction, consistent with reward encoding of shock withdrawal. Conversely, silencing PAM-β’2a by activating GtACR1 light-gated inhibitory anion channel (Mohammad et al., 2017) with green light during CS+ presentation in reversal trials increases CS+ avoidance relative to control, reflecting impairment of extinction of the aversive CS+ association (**Figure 1J**). This indicates that PAM-β’2a activity during CS+ presentation in reversal trials is necessary to drive extinction of the aversive CS+ association, again consistent with reward encoding of shock withdrawal. Importantly, PAM-β’2a does not encode a general reward signal, as optogenetic activation of PAM-β’2a during *de novo* odor presentation is insufficient to generate a positive association (**Figure S1E**), and optogenetic inhibition fails to interfere with aversive shock-odor memory acquisition (**Figure S1F**). Taken together, these imaging and functional optogenetic results indicate a specific role for PAM-β’2a in encoding the reward of punishment withdrawal during reversal.

### PAM-β’2a decreases CS+ odor avoidance during reversal by depressing KC-to-MBON-γ5β’2a synapses

How does the increase in PAM-β’2a CS+ odor response during reversal decrease CS+ avoidance? Since PAM-β’2a innervates the same MB compartments as MBON-γ5β’2a and MBON-β2β’2a (Aso et al., 2014a), we focused our attention on these two MBONs, which encode negative and positive valence respectively (Aso et al., 2014b). MBON-γ5β’2a increases its CS+ odor response during acquisition (Bouzaiane et al., 2015; Owald et al., 2015; Felsenberg et al., 2018) and decreases CS+ odor response during reversal (**Figure 2A**,**B**), consistent with previous studies (Bouzaiane et al., 2015; Owald et al., 2015; Felsenberg et al., 2018). These changes do not occur during mock conditioning (**Figure 2C**). MBON-β2β’2a increases its response to both CS+ and CS-odors during acquisition, and no changes in odor response are observed during reversal (**Figure S2A**). These imaging results suggest that MBON-γ5β’2a is the downstream target through which PAM-β’2a reduces CS+ avoidance during reversal, as reduced MBON-γ5β’2a CS+ odor response would encode reduced negative valence of the CS+ odor.

**Figure 2:**
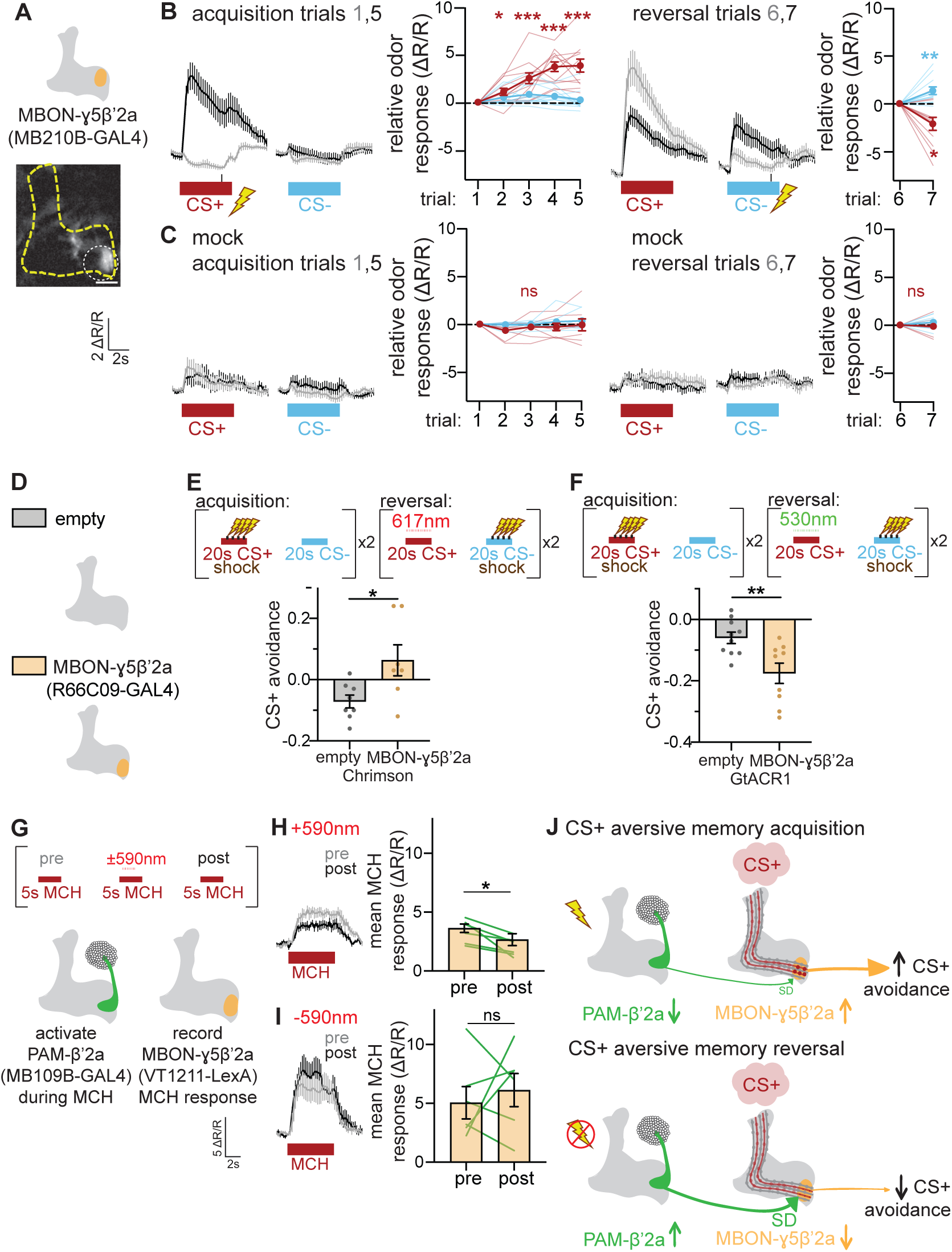
PAM-β’2a DANs induce depression of KC-MBON-γ5β’2a synapses to decrease CS+ odor avoidance during reversal. (**A**) Schematic and sample fluorescence image of MBON-γ5β’2a, an output neuron receiving dopaminergic input from PAM-β’2a. (**B**) Neural activity of MBON-γ5β’2a in flies undergoing aversive memory acquisition and reversal. MBON-γ5β’2a CS+ odor response increases during acquisition and decreases during reversal. *n* = 12 flies. (**C**) Neural activity of MBON-γ5β’2a in flies undergoing mock aversive memory acquisition and reversal. No change in odor response occurs during acquisition or reversal. *n* = 8 flies. (**D**) Schematic of MBON-γ5β’2a and empty-splitGAL4 control used in behavioral experiments. (**E**) Activating MBON-γ5β’2a during CS+ delivery in reversal impairs extinction of CS+ aversive memory, reflected in increased CS+ avoidance. *n* = 8 & 7 independent groups of flies per genotype. (**F**) Silencing MBON-γ5β’2a during CS+ delivery in reversal enhances extinction, reflected in decreased CS+ avoidance. *n* = 10 & 9 independent groups of flies per genotype. (**G**) Top: Optogenetic manipulation of DAN-mediated synaptic plasticity. Odor (MCH) is presented to naïve flies with 3s pulse of red light to activate Chrimson in PAM-β’2a. Odor-evoked MBON-γ5β’2a neural responses to MCH are recorded 2 minutes before (pre) and after (post) odor-light pairing. Bottom: Flies express red-light-activated Chrimson in PAM-β’2a neurons and GCaMP6m in MBON-γ5β’2a to stimulate and visualize neural activity, respectively. (**H**) Neural responses in MBON-γ5β’2a before (grey) and after (black) odor-light pairing. Optogenetic activation of PAM-β’2a during MCH odor delivery decreases subsequent MCH odor response. *n* = 6 flies. Statistical comparison is by paired *t*-test. (**I**) Same experiment as above, except mock trials without red light illumination. Odor responses in MBON-γ5β’2a before (grey) and after (black) mock odor-light pairing are not significantly different. *n* = 6 flies. Statistical comparison is by paired *t*-test. (**J**) Schematized circuit mechanisms involving PAM-β’2a and MBON-γ5β’2a during aversive memory acquisition and reversal, using a pink cloud and a lightning bolt to depict CS+ odor and electric shock, respectively. Mushroom body is depicted in grey; parallel fibers and synapses represent lines and circles, respectively. Maroon lines and circles indicate KC parallel fibers activated by CS+ odor. Size of circles indicate strength of KC-MBON synapses. Direction of arrowhead next to each neuron’s name indicates direction of change in CS+ odor response (i.e., increase or decrease). Thickness of arrow and size of arrowhead indicate said changes in CS+ odor response; they additionally show the direction of information flow in the circuit. For clarity, only one hemisphere is depicted, and DANs and MBONs are depicted on the left and right side respectively. During acquisition, CS+ odor response of PAM-β’2a (green) decreases, reducing any synaptic depression (SD) induced onto KC-MBON-γ5β’2a synapses. (Feedforward disinhibition of MBON-γ5β’2a CS+ odor response via the γ1ped compartment known to occur during acquisition is not depicted.) During reversal, CS+ odor is presented without electric shock. The unexpected withdrawal of electric shock increases PAM-β’2a CS+ odor response. This induces synaptic depression at the KC-MBON-γ5β’2a synapse (small red circles), thus decreasing MBON-γ5β’2a (orange) CS+ odor response and thereby reducing CS+ avoidance.

We optogenetically interrogated whether avoidance-encoding MBON-γ5β’2a is involved in reversal. Optogenetic activation of MBON-γ5β’2a during CS+ presentation in reversal trials impairs the extinction of the aversive CS+ association, reflected in relative increase in CS+ avoidance compared to control (**Figure 2E**). This likely occurs by counteracting the decrease in CS+ odor response that normally occurs during reversal (see **Figure 2B**). Conversely, optogenetic inhibition of MBON-γ5β’2a enhanced extinction of the aversive CS+ association, reflected in a relative decrease in CS+ odor avoidance (**Figure 2F**). These results indicate that avoidance-encoding MBON-γ5β’2a plays a causal role in decreasing CS+ avoidance during reversal.

To test whether PAM-β’2a modulates MBON-γ5β’2a odor responses, we employed a combination of optogenetics and Ca^2+^ imaging. *De novo* odor presentation of 4-methyl-cyclohexanol (“MCH”) is paired with a 3s pulse of red light to activate Chrimson-expressing PAM-β’2a in naïve flies, and odor-evoked responses are measured in GCaMP6m-expressing MBON-γ5β’2a before and after the odor-light pairing (**Figure 2G**). PAM-β’2a activation during odor presentation decreases subsequent odor responses of MBON-γ5β’2a, consistent with sustained synaptic depression of KC-MBON-γ5β’2a synapses (**Figure 2H**). MBON-γ5β’2a odor responses do not change in independent mock experiments where no light is delivered (**Figure 2I**). No effects of odor-light pairing on subsequent MBON-γ5β’2a odor responses occur in empty-splitGAL4 control flies (**Figure S2C**). We also quantified odor responses in MBON-β’2mp, which are unaffected by PAM-β’2a activation during prior odor presentation (**Figure S2B**). These results demonstrate that PAM-β’2a activation during odor presentation depresses KC-MBON-γ5β’2a synapses persistently, thus decreasing MBON-γ5β’2a odor response. This is consistent with the previous observation that pairing neutral odor with broad, non-specific activation of PAMs decreases MBON-γ5β’2a odor response (Handler et al., 2019).

Taken together, the experiments presented thus far support a cellular and synaptic mechanism for CS+ extinction during reversal learning (**Figure 2J**): During acquisition, PAM-β’2a decreases CS+ odor response. When electric shock is withdrawn during reversal, PAM-β’2a increases CS+ odor response, thus depressing KC-MBON-γ5β’2a synapses, decreasing MBON-γ5β’2a CS+ odor response, and thereby decreasing CS+ avoidance.

### MBON-γ2α’1 is upstream of PAM-β’2a and drives CS+ extinction induced by shock withdrawal

What causes the change in PAM-β’2a CS+ odor response during acquisition and reversal? Since DAN dendrites receive extensive inputs from MBONs (Aso et al., 2014a), we again took an unbiased screening approach to identify MBONs in addition to MBON-γ5β’2a that change their CS+ odor response during reversal learning.

Out of 17 MBON subsets screened, MBON-γ1ped and MBON-γ5β’2a changed CS+ odor response during acquisition (**Figure 3A**), consistent with prior observations (Bouzaiane et al., 2015; Hige et al., 2015; Owald et al., 2015; Perisse et al., 2016). MBON-α’2, MBON-β’2mp, MBON-γ1ped, MBON-γ2α’1 and MBON-γ5β’2a changed CS+ odor response relative to CS-during reversal (**Figure 3B**). MBON-α’2 and MBON-β’2mp increase CS+ odor response during acquisition, and decrease during reversal (**Figure 3C**,**D**). Increased CS+ odor response of MBON-β’2mp during acquisition has been observed previously (Owald et al., 2015). MBON-γ1ped decreases CS-odor response during reversal, while its CS+ odor response is unchanged (**Figure 3E**), also consistent with prior observations (Felsenberg et al., 2018). In contrast to MBON-α’2 and MBON-β’2mp, MBON-γ2α’1 decreases CS+ odor response during acquisition and increases during reversal (**Figure 3F**), consistent with prior observations (Berry et al., 2018). No changes in CS+ odor response of MBON-β’2mp, MBON-γ1ped or MBON-γ2α’1 occur during mock reversal learning without shock (**Figure S3A-C**). These changes in MBON-γ1ped and MBON-γ2α’1 CS+ odor responses also occur when the odors used as CS+ and CS-are switched (**Figure S3B**,**C**).

**Figure 3:**
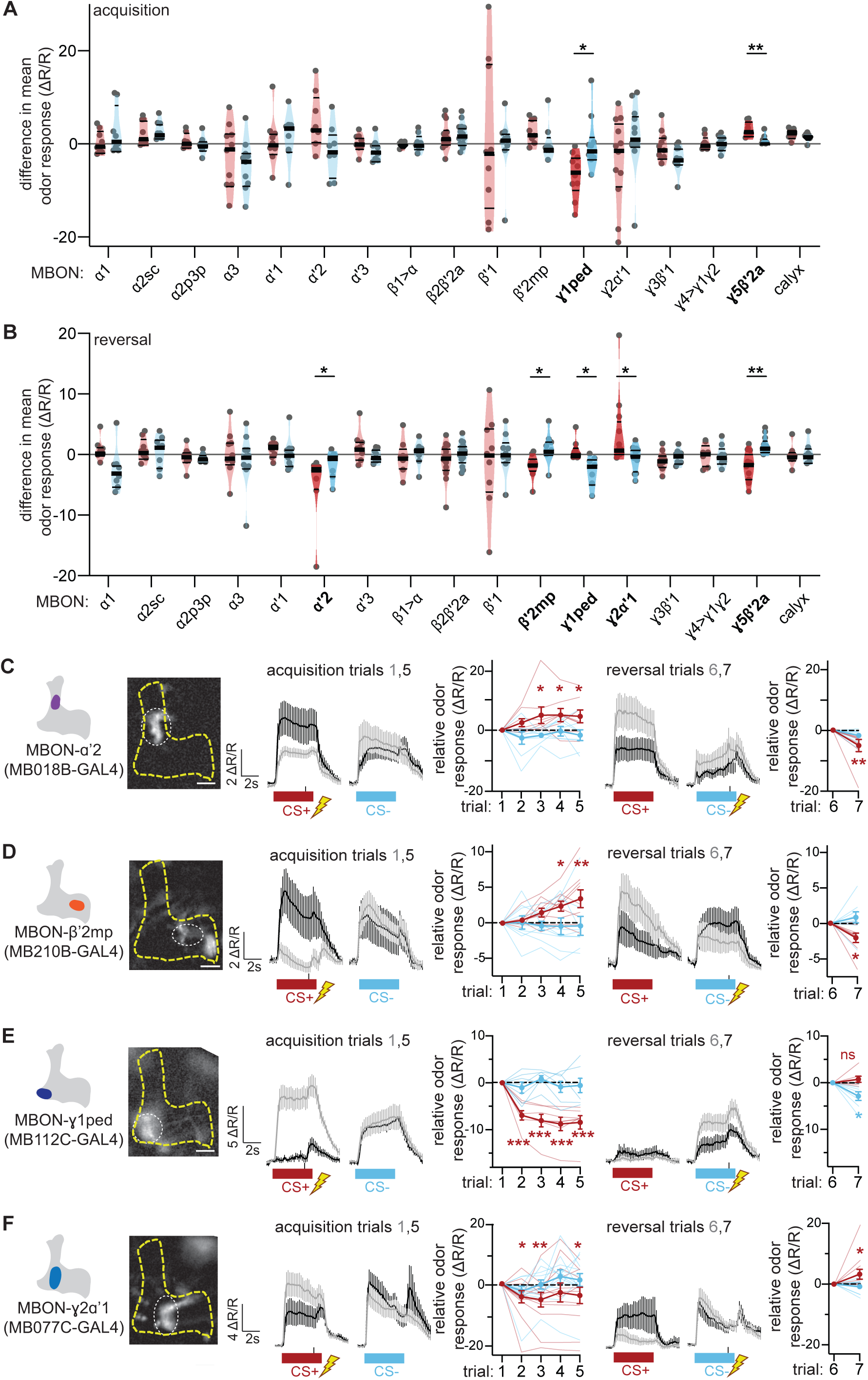
Five MBONs significantly change odor responses during reversal. (**A**) MBON imaging screen: acquisition trials. Relative difference in mean Ca^2+^ response of indicated MBONs to CS+ and CS-odors from last to first acquisition trials. Change in MBON-γ1ped and MBON-γ5β’2a CS+ odor response relative to CS-odor responses are significantly different. *n* = 7-14 flies per genotype. Statistical comparison is by paired *t*-test or Wilcoxon matched-pairs signed rank test between CS+ and CS-odor response for each genotype. (**B**) MBON imaging screen: reversal trials. Changes in CS+ odor responses relative to CS-odor responses in five MBONs – α’2, β’2mp, γ1ped, γ2α’1 and γ5β’2a – are significantly different. (**C**) MBON-α’2 CS+ odor response increases during acquisition and decreases during reversal. *N* = 8 flies. (**D**) MBON-β’2mp CS+ odor response increases during acquisition and decreases during reversal. *n* = 9 flies. (**E**) MBON-γ1ped CS+ odor response decreases during acquisition, and its CS-odor response decreases during reversal. *n* = 7 flies. Since its CS+ odor response does not change during reversal, it is excluded as a possible upstream modulator of PAM-β’2a. (**F**) MBON-γ2α’1 CS+ odor response decreases during acquisition and increases during reversal. *n* = 12 flies.

MBON-α’2 does not appear to encode intrinsic valence (Aso et al., 2014b), nor is there any obvious anatomical substrate for connectivity with PAM-β’2a (Aso et al., 2014a). We thus optogenetically interrogated MBON-γ2α’1 and MBON-β’2mp as potential upstream partners of PAM-β’2a that could encode shock withdrawal during reversal (**Figure 4A**). Activating Chrimson-expressing MBON-γ2α’1 with maximal red light exposure during CS+ delivery in reversal trials enhances extinction of the aversive CS+ memory, reflected in decreased CS+ avoidance relative to empty-splitGAL4 control during subsequent preference testing (**Figure 4B**). In contrast, activating MBON-β’2mp with lowest red light exposure impairs extinction, reflected in increased CS+ avoidance relative to control (**Figure 4B**). These effects during reversal parallel previously reported rewarding and punishing intrinsic valences encoded by MBON-γ2α’1 and MBON-β’2mp, respectively (Aso et al., 2014b). Optogenetic inhibition of GtACR1-expressing MBON-γ2α’1 or MBON-β’2mp with lowest green light exposure during CS+ delivery in reversal trials impairs and enhances extinction, respectively (**Figure 4C**). These effects of optogenetic manipulation of MBON-γ2α’1 and MBON-β’2mp are again consistent with roles for both of these neurons in encoding shock withdrawal during CS+ delivery in reversal trials.

**Figure 4:**
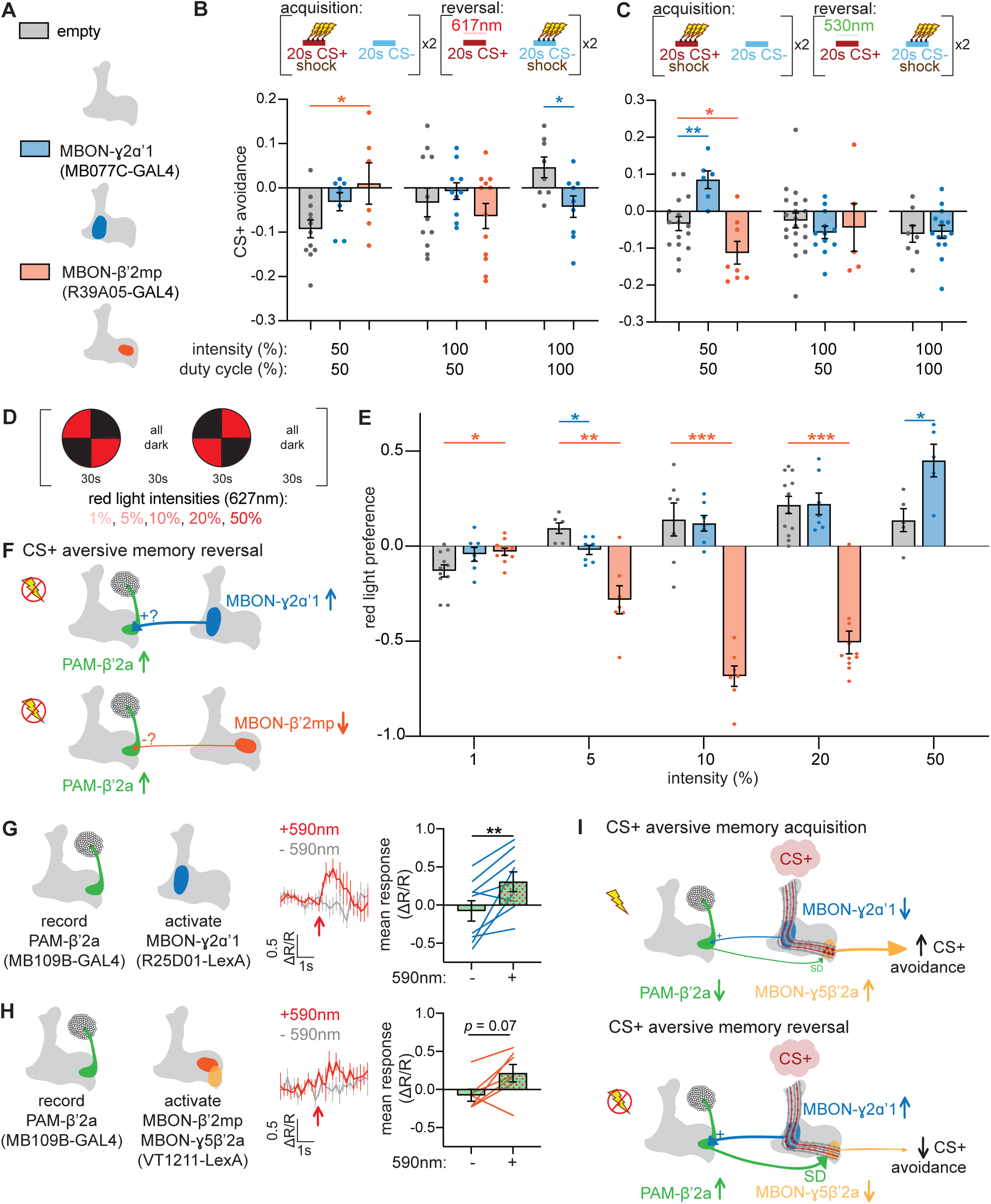
MBON-γ2α’1 activation of PAM-β’2a encodes shock withdrawal during reversal learning. (**A**) Schematic of MBON-γ2α’1, MBON-β’2mp and empty-splitGAL4 control driver lines. (**B**) Activating approach-encoding MBON-γ2α’1 during CS+ reversal with the highest light dose enhances extinction of CS+ aversive memory, reflected in decreased in CS+ avoidance relative to control. Activating avoidance-encoding MBON-β’2mp during CS+ reversal with the lowest light dose impairs extinction of CS+ aversive memory, reflected in increased in CS+ avoidance relative to control. *n* = 6-12 independent groups of flies per genotype. Statistical comparison is by two-way ANOVA with Dunnett’s multiple comparisons test against empty-splitGAL4 control. (**C**) Silencing approach-encoding MBON-γ2α’1 during CS+ reversal at the lowest light dose impairs extinction of CS+ aversive memory, reflected in increased CS+ avoidance relative to control. Silencing avoidance-encoding MBON-β’2mp during CS+ reversal at the lowest light dose enhances extinction of CS+ aversive memory, reflected in decreased CS+ avoidance relative to control. *n* = 5-20 independent groups of flies per genotype. Statistical comparison is by two-way ANOVA with Dunnett’s multiple comparisons test against empty-splitGAL4 control. (**D**) Experimental protocol for assessing innate valence encoded by each MBON. About 30 naïve flies are placed in a dark circular arena and allowed to move freely. After one minute of acclimation, two quadrants are illuminated with red light for 30 seconds to activate Chrimson-expressing neurons. After 30 seconds, the other two quadrants are illuminated. The preference for red light quadrants is averaged to calculate the final preference score for that particular red light intensity. This is then repeated using light of increasing intensities. (**E**) A red light preference value of 1 indicates that 100% of the flies prefer the red light quadrants; a preference of 0 indicates that flies are indifferent to lit versus non-lit quadrants. Control flies (empty, grey) avoid red-light-illuminated quadrants at the lowest light intensity, and increase their preference for red-light-illuminated quadrants at higher intensities. Flies expressing Chrimson in MBON-β’2mp are relatively attracted to red light quadrants at the lowest light intensity, and avoid red light quadrants relative to control at higher intensities. Flies expressing Chrimson in MBON-γ2α’1 avoid red light quadrants relative to control at low light intensity, and prefer red-light quadrants relative to control at the highest light intensity. *n* = 4-13 groups of flies per genotype and condition. Statistical comparison is by two-way ANOVA with Dunnett’s multiple comparisons test against empty-splitGAL4 control. (**F**) Proposed connectivity between potential upstream neurons and PAM-β’2a. Since MBON-γ2α’1 and MBON-β’2mp increase and decrease their CS+ odor response during reversal, respectively, they would have to excite (+) and inhibit (-) PAM-β’2a respectively if they are the upstream neuron involved, since PAM-β’2a increases CS+ odor response during reversal. (**G**) Chrimson-expressing MBON-γ2α’1 is activated with 500ms pulse of red light, and neural response in GCaMP6m-expressing PAM-β’2a is recorded. Neural activity is first recorded without stimulation (grey); five minutes later, neural activity is recorded with red light stimulation occurring 10s after the imaging lights are turned on (red). Red arrow indicates onset of red light. Right: quantification of mean neural activity 1s after red light onset. Activating MBON-γ2α’1 increases PAM-β’2a activity. *n* = 9 flies. Statistical comparison is by paired *t*-test. (**H**) Activating MBON-β’2mp and MBON-γ5β’2a does not cause a significant change in PAM- β’2a activity. *n* = 7 flies. Statistical comparison is by Wilcoxon signed-rank test. (**I**) Proposed role of MBON-γ2α’1 and PAM-β’2a during aversive memory acquisition and reversal. During memory acquisition, MBON-γ2α’1 CS+ odor response decreases, leading to a decrease in PAM-β’2a CS+ odor response via an excitatory connection (+). During reversal, the unexpected withdrawal of electric shock restores MBON-γ2α’1 CS+ odor response and increases PAM-β’2a CS+ odor response. This drives synaptic depression at the KC-MBON-γ5β’2a synapse, decreasing MBON-γ5β’2a odor response, which reduces CS+ avoidance.

The complex relationship between degree of light exposure and behavioral effects in these optogenetic experiments could be due to (1) photobiophysics and membrane biophysics of Chrimson and GtACR1-expressed in MBON-γ2α’1 and MBON-β’2mp, (2) nonlinear network effects in the context of recurrent MB connectivity (Aso et al., 2014a), and/or (3) superimposed non-linear intrinsic effects of red and green light on behavior. We assessed the effect of red light intensity on approach/avoidance to the light itself by flies expressing Chrimson in MBON-γ2α’1, MBON-β’2mp, and non-expressing empty-splitGAL4 control flies (**Figure 4D**). Control flies avoid the lowest intensity red light, and shift to attraction at higher intensities. Flies expressing Chrimson in MBON-γ2α’1 shift from relative avoidance to relative attraction compared to control as light intensity increases (**Figure 4E**). Conversely, flies expressing Chrimson in MBON-β’2mp shift from relative attraction to relative avoidance as light intensity increases (**Figure 4E**).

These intensity-dependent differences in approach/avoidance to light in some respects parallel the intensity-dependent effects of light in the context of reversal learning (**Figure 4B**,**C**). For example, activation of Chrimson-expressing MBON-γ2α’1 with higher red light dose during CS+ delivery in reversal trials decreases CS+ avoidance relative to control during subsequent testing (**Figure 4B**), while lower doses of red light do not. Similarly, only the highest red light intensity drives approach of flies expressing Chrimson in MBON-γ2α’1, while lower intensities do not (**Figure 4E**). For flies expressing Chrimson in MBON-β’2mp, only the lowest red light dose during CS+ delivery in reversal trials increases CS+ avoidance during subsequent testing, and not a higher dose (**Figure 4B**). Similarly, red light preference of flies expressing Chrimson in MBON-β’2mp differs between the lowest and higher light intensities (**Figure 4E**).

To distinguish potential roles of MBON-γ2α’1 and MBON-β’2mp in conveying withdrawal of shock during CS+ reversal trials to PAM-β’2a, we optogenetically activated each of these neurons individually using Chrimson while recording neural activity of PAM-β’2a. Since MBON-γ2α’1 and MBON-β’2mp increase and decrease CS+ odor response during reversal, respectively, they would need to excite and inhibit PAM-β’2a, respectively, if involved in increasing PAM-β’2a CS+ odor response during reversal (**Figure 4F**). Optogenetic activation of MBON-γ2α’1 excites PAM-β’2a (**Figure 4G**), supporting MBON-γ2α’1 as transmitting information of withdrawal of shock during CS+ presentation in reversal trials to PAM-β’2a. This is consistent with the prior observation that activating MBON-γ2α’1 excites PAMs, although use of a broad driver for GCaMP expression precluded subset identification (Felsenberg et al., 2017). Red light fails to elicit PAM-β’2a responses in control flies that do not express Chrimson (**Figure S4A**). Optogenetic activation of MBON-β’2mp and MBON-γ5β’2a (no MBON-β’2mp specific LexA driver exists) fails to elicit PAM-β’2a responses, and particularly, fails to inhibit PAM-β’2a (**Figure 4H**), as would be required for an upstream neuron whose CS+ odor response decreases during reversal to transmit withdrawal of shock information to reward-encoding PAM-β’2a.

Taken together, the experiments presented thus far support the following model (**Figure 4I**): During acquisition, decreased MBON-γ2α’1 CS+ odor response decreases excitation of PAM-β’2a, reflected in its decreased CS+ odor response. During reversal, withdrawal of shock increases MBON-γ2α’1 CS+ odor response, thus increasing PAM-β’2a CS+ odor response. This increases persistent synaptic depression of KC-MBON-γ5β’2a synapses, thus persistently decreasing MBON-γ5β’2a CS+ odor response and, consequently, decreasing CS+ odor avoidance during subsequent testing.

### Shock-responsive PPL1-γ2α’1 conveys shock withdrawal to avoidance-encoding MBON-γ5β’2a through MBON-γ2α’1-to-PAM-β’2a relay

Shock-responsive PPL1-γ2α’1 is the only DAN that projects its dopaminergic terminals to the same MB compartment innervated by the dendrites of MBON-γ2α’1 (Berry et al., 2018). This leads to the hypothesis that shock withdrawal information enters the circuit via PPL1-γ2α’1, and is relayed to avoidance-encoding MBON-γ5β’2a via MBON-γ2α’1 and PAM-β’2a.

To test this hypothesis, we optogenetically interrogated whether reduced PPL1-γ2α’1 activity during shock withdrawal is required for reversal learning. Substitution of direct optogenetic activation of PPL1-γ2α’1 for shock during CS+ presentation in reversal trials prevents CS+ extinction in flies expressing Chrimson in PPL1-γ2α’1, reflected in increased CS+ avoidance compared to control flies not expressing Chrimson (**Figure 5A**). Optogenetic shock substitution targeted to other shock-responsive DAN subsets (**Figure S5A**,**B**) presynaptic to other MBONs that changed CS+ odor response during reversal (**Figure 3**) fails to prevent CS+ extinction (**Figure 5A**).

**Figure 5:**
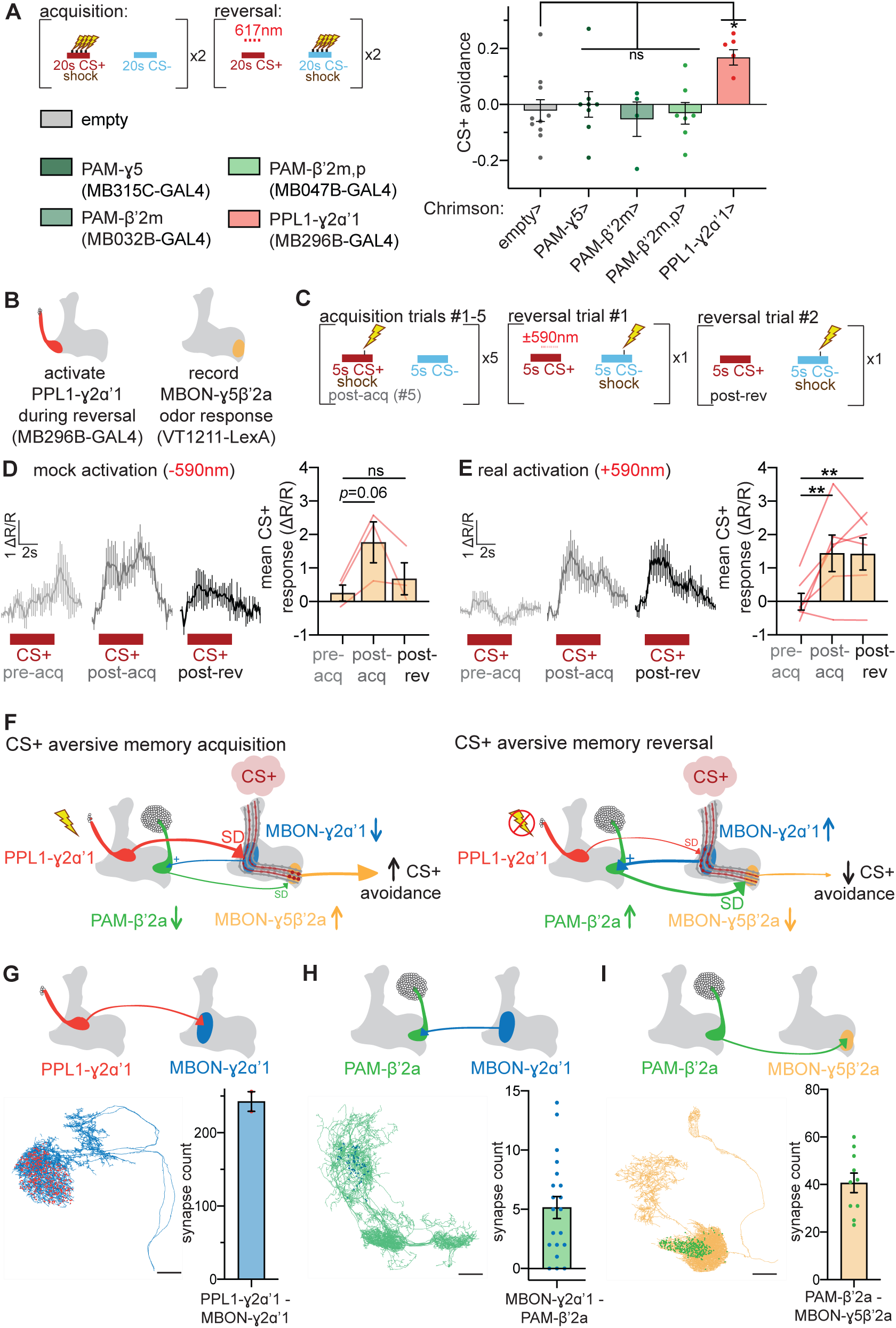
Shock-responsive PPL1-γ2α’1 dopaminergic neurons convey withdrawal of shock during reversal. (**A**) Experimental paradigm for “shock-substitution” experiment, in which pulses of red light are used to mimic electric shock during reversal. When Chrimson is expressed in PPL1-γ2α’1, red light pulses delivered during CS+ reversal trials impair reversal, reflected in the relative increase in CS+ avoidance. In contrast, activation of other shock-responsive DANs during CS+ delivery fails to impair reversal. *n* = 10, 8, 4, 7, & 5 independent groups of flies per genotype. Statistical comparison is by one-way ANOVA with Dunnett’s post-hoc test. (**B**) Flies express red-light-activated Chrimson in PPL1-γ2α’1 neurons, and GCaMP6m in MBON-γ5β’2a. (**C**) Experimental paradigm for determining effect of PPL1-γ2α’1 activation on reversal-induced decrease in MBON-γ5β’2a CS+ odor response. During pre-acquisition, CS+ and CS-odors are alternately presented to the fly once, and MBON-γ5β’2a CS+ odor response is recorded (“pre-acq”). For five acquisition trials, CS+ is paired with electric shock and the CS-is presented without shock. On the 5^th^ acquisition trial, CS+ odor response is recorded (“post-acq”). During the first reversal trial, CS+ is presented with a 3s pulse of red light to activate PPL1-γ2α’1. On second reversal trial, CS+ odor response is recorded (“post-rev”). (**D**) Neural responses in MBON-γ5β’2a without red-light activation of PPL1-γ2α’1. MBON-γ5β’2a increases and decreases CS+ odor response during acquisition and reversal respectively. *n* = 3 flies. Statistical comparison is by one-way ANOVA with Dunnett’s post-hoc test against pre-acquisition. (**E**) Optogenetic activation of PPL1-γ2α’1 DAN during CS+ presentation in first reversal trial prevents decrease in MBON-γ5β’2a CS+ odor response. *n* = 6 flies. Statistical comparison is by one-way ANOVA with Dunnett’s post-hoc test against pre-acquisition. (**F**) Mechanism of aversive memory acquisition and reversal. During memory acquisition, coincident activation of specific KCs by the CS+ odor and PPL1-γ2α’1 by electric shock causes synaptic depression at the KC-MBON-γ2α’1 synapse, leading to a decrease in MBON-γ2α’1 CS+ odor response. This causes a decrease in PAM-β’2a CS+ response, which reduces synaptic depression induced on the KC-MBON-γ5β’2a synapse, thus potentiating the increase in CS+ odor response by MBON-γ5β’2a. During reversal, the withdrawal of electric shock, coupled by the pairing of electric shock with the CS-odor, restores plasticity at the KC-MBON-γ2α’1 synapse. The increase in MBON-γ2α’1 CS+ odor response excites PAM-β’2a, leading to its increased CS+ odor response. This induces synaptic depression on its postsynaptic MBON, leading to decreased MBON-γ5β’2a CS+ odor response and a decrease in CS+ odor avoidance. (**G**) Synapses between PPL1-γ2α’1 and MBON-γ2α’1. Left, blue neuron is EM-traced MBON-γ2α’1; red dots represent location of individual synapses from PPL1-γ2α’1. Right: quantification of number of synapses from PPL1-γ2α’1 onto each MBON-γ2α’1 neuron. Several synapses are formed between PPL1-γ2α’1 and the dendritic regions of both MBON-γ2α’1 neurons, making PPL1-γ2α’1 a likely modulator of KC-MBON-γ2α’1 synapses. *n* = 2, for the two MBON-γ2α’1 neurons. Scale bar is 20μm. (**H**) Synapses between MBON-γ2α’1 and PAM-β’2a. Green neuron is EM-traced PAM-β’2a neurons; blue dots represent individual synapses from MBON-γ2α’1. Several synapses exist between the two MBON-γ2α’1 neurons and the dendritic region of ten PAM-β’2a neurons, making the excitatory connection between MBON-γ2α’1 and PAM-β’2a identified in **Figure 4** likely a monosynaptic one. *n* = 20, for the connections between two MBON-γ2α’1 neurons and ten PAM-β’2a neurons. Scale bar is 20μm. (**I**) Synapses between PAM-β’2a and MBON-γ5β’2a. Orange neuron is EM-traced MBON-γ5β’2a; green dots represent individual synapses from PAM-β’2a. Several synapses exist between each of the ten PAM-β’2a neurons and the dendritic regions of MBON-γ5β’2a, making PAM-β’2a a likely modulator of KC-MBON-γ5β’2a synapses. *n* = 10, for the connections between ten PAM-β’2a neurons and MBON-γ5β’2a. Scale bar is 20μm.

We also combined functional imaging and optogenetics by activating Chrimson-expressing PPL1-γ2α’1 during CS+ delivery in reversal trial #1, and then measuring CS+ odor response of GCaMP6m-expressing MBON-γ5β’2a during reversal trial #2 (**Figure 5B**,**C**). In mock experiments in which no red light is presented, CS+ odor responses of MBON-γ5β’2a increase after acquisition and then decrease after reversal (**Figure 5D**), as in **Figure 2B**. In contrast, when Chrimson-expressing PPL1-γ2α’1 is activated with red light during CS+ odor delivery in reversal trial #1, the decrease in CS+ odor response in reversal trial #2 is eliminated (**Figure 5E**). This indicates that encoding of withdrawal of shock by PPL1-γ2α’1 is necessary for MBON-γ5β’2a CS+ odor response to decrease during reversal. Previous studies reveal that, like shock itself, direct optogenetic activation of PPL1-γ2α’1 during odor presentation changes subsequent MBON-γ2α’1 odor responses (Cohn et al., 2015; Berry et al., 2018). Our results indicate that PPL1-γ2α’1 encoding of shock is uniquely necessary during reversal learning to convey shock withdrawal to avoidance-encoding MBON-γ5β’2a via an MBON-γ2α’1-to-PAM-β’2a indirect dopaminergic relay (**Figure 5F**).

To assess synaptic connectivity potentially underlying the circuit we have presented multiple lines of functional evidence for, we interrogated the recently published *Drosophila* hemibrain electron microscopy volume (Scheffer et al., 2020; Xu et al., 2020). There are numerous synaptic connections from the single PPL1-γ2α’1 DAN onto each of the two MBON-γ2α’1 neurons (**Figure 5G**). Each of the two MBON-γ2α’1 neurons forms an average of five synaptic connections onto the dendrites of each of the ten PAM-β’2a DANs (**Figure 5H**). MBON-γ1ped, MBON-γ3β’1, MBON-γ4γ5, and MBON-γ5β’2a each form substantially fewer synaptic connections onto PAM-β’2a than MBON-γ2α’1 (**Figure S6A**). Each PAM-β’2a neuron forms dozens of synaptic connections onto MBON-γ5β’2a (**Figure 5I**). These synaptic connections visualized directly via electron microscopy are consistent with our model, and suggest that the functional connections we have uncovered optogenetically are monosynaptic.

## DISCUSSION

Here we combine functional imaging and optogenetic interrogation to provide extensive evidence for a lateral relay connecting the γ2α’1 MB microcircuit to the γ5β’2a microcircuit to encode shock withdrawal during reversal learning as reward. While dopaminergic neurons have been generally implicated in encoding punishment withdrawal as reward, for the first time we identify and dissect synaptic connectivity and modulatory roles in synaptic plasticity of specific DANs encoding shock withdrawal as reward.

Our results support the following model for how shock withdrawal extinguishes the CS+-shock association and reduces CS+ avoidance: During aversive memory acquisition, coincident activation of specific KCs by the CS+ odor and PPL1-γ2α’1 by electric shock depresses KC-MBON-γ2α’1 synapses, decreasing MBON-γ2α’1 CS+ odor response (Berry et al., 2018). This decreases PAM-β’2a CS+ response, thus relieving synaptic depression of KC-MBON-γ5β’2a synapses, and increasing MBON-γ5β’2a CS+ odor response. This PAM-β’2a-mediated increase in MBON-γ5β’2a CS+ odor response is in addition to the known increase in MBON-γ5β’2a CS+ odor response caused by disinhibition of MBON-γ1ped (Hige et al., 2015; Perisse et al., 2016; Felsenberg et al., 2018). During reversal, the withdrawal of electric shock during CS+ odor presentation decreases PPL1-γ2α’1 activation, which relieves depression of KC-MBON-γ2α’1 synapses. Consequent increase in MBON-γ2α’1 CS+ odor response is relayed by an excitatory synapse to PAM-β’2a, increasing its CS+ odor response. This depresses KC-MBON-γ5β’2a synapses, decreases MBON-γ5β’2a CS+ odor response, and thereby decreases CS+ odor avoidance. Thus, synaptic plasticity in the γ2α’1 microcircuit directly responds to shock withdrawal, and conveys this information to the γ5β’2a microcircuit via PAM-β’2a to extinguish the CS+-shock association.

While PAM-β’2a has not previously been shown to encode shock withdrawal as reward, it has been previously implicated as encoding other rewards in other contexts, such as water-seeking (Lin et al., 2014), sugar-mediated appetitive learning (Huetteroth et al., 2015), water (Senapati et al., 2019) or ethanol (Scaplen et al., 2019) reinforcement, and food-mediated suppression of CO2 avoidance (Lewis et al., 2015). In the case of ethanol reinforcement, PAM-β’2a likely acts through MBON-β2β’2a, not MBON-γ5β’2a (Scaplen et al., 2019), indicating that PAM-β’2a conveys reward information in different contexts via different downstream MBONs.

The involvement of avoidance-encoding MBON-γ5β’2a in reversal learning is consistent with its similar role in extinction learning (Felsenberg et al., 2018). In both cases, MBON-γ5β’2a CS+ odor response increases during acquisition and decreases when CS+ is no longer presented with shock. The authors speculated that the increase in MBON-γ5β’2a CS+ odor response during acquisition causes an increase in PAM-γ5 CS+ odor response, which then induces synaptic depression at KC-MBON-γ5β’2a synapses to reduce MBON-γ5β’2a CS+ odor response. However, it is not clear how this would occur specifically during extinction and not acquisition, since no circuit encoding shock information is proposed. Furthermore, neither they nor we found changes in PAM-γ5 CS+ odor response during acquisition or extinction/reversal. It would be interesting to see whether PAM-β’2a is also the cause of the decrease in MBON-γ5β’2a CS+ odor response during extinction learning, or whether its modulation of plasticity indeed comes from PAM-γ5 as they speculate (Felsenberg et al., 2018; Otto et al., 2020). Unlike MBON-γ5β’2a, some MBONs (e.g., MBON-γ2α’1 and MBON-β’2mp) change their CS+ odor response during extinction learning and not reversal learning, so it would not be surprising if non-overlapping DAN populations are involved in different tasks.

Anatomical studies reveal both intra- and inter-compartment connections between DANs and MBONs (Aso et al., 2014a; Takemura et al., 2017), and functional studies suggest the involvement of inter-compartment connections in some forms of learning (Cognigni et al., 2018; Eschbach et al., 2019; Modi et al., 2020). MBON-γ2α’1 makes connections with a variety of DAN subsets, and is involved in appetitive and aversive short-term memory (Berry et al., 2018; Yamazaki et al., 2018) and consolidation of appetitive memories (Felsenberg et al., 2017). PPL1-γ2α’1 has also been implicated in acquisition of appetitive memories (Berry et al., 2018; Yamazaki et al., 2018), the acquisition and forgetting of aversive memories (Aso et al., 2012; Berry et al., 2012; Placais et al., 2012; Berry et al., 2015; Aso and Rubin, 2016) and reconsolidation of CS-memory encoding absence of reward (Felsenberg et al., 2017). Here we demonstrate a role for MBON-γ2α’1 and PPL1-γ2α’1 in encoding shock withdrawal as reward during reversal learning.

We propose that the γ1ped microcircuit encoding the originally acquired aversive memory (Hige et al., 2015; Perisse et al., 2016; Felsenberg et al., 2018), and the γ2α’1-β’2 relay network we uncover here as encoding a newly formed reward memory, are analogous to the roles of the amygdala and ventromedial prefrontal cortex (vmPFC) networks in fear reversal learning in mammals. The amygdala persistently encodes the originally acquired fear memory (Schiller et al., 2008; Schiller and Delgado, 2010; Li et al., 2011), while the vmPFC encodes omission of the aversive outcome during reversal as reward (Schiller et al., 2008; Zhang et al., 2015). Interestingly, inhibiting vmPFC-projecting mesocortical ventral tegmental area (VTA) DANs during shock omission impairs extinction (Luo et al., 2018), analogous to our finding that inhibiting PAM-β’2a CS+ odor response during reversal impairs extinction of the aversive memory. PAM-β’2a odor responses can thus be thought of as a prediction error signal, which encode the value associated with each cue by iteratively updating the value on a trial-by-trial basis (Rescorla, 1972; Schultz et al., 1997). Since a subset of VTA DANs have been found to respond to aversive stimuli such as electric shock (de Jong et al., 2019), it would be interesting to see if these aversive-encoding DANs relay information about absence or presence of shock to reward-encoding DANs which signal unexpected withdrawal of shock to the vmPFC. In sum, we have revealed how punishment-encoding DANs and reward-encoding DANs collaborate in an indirect dopaminergic relay circuit in the fly to encode shock withdrawal as reward, and thereby underlie formation of a CS+ reward memory during reversal learning. This circuit topology for encoding withdrawal of punishment as reward could be conserved across animal species, and suggests testable hypotheses for exploration in mammals.

## EXPERIMENTAL PROCEDURES

### Fly husbandry

Fly stocks were cultured on standard cornmeal medium at 25°C with a 12hr light-dark cycle. Flies used for functional connectivity experiments were maintained on molasses medium. Flies used for optogenetic behavioral or functional connectivity experiments were kept in the dark using aluminum foil, and raised on standard cornmeal medium supplemented with 0.4mM all-*trans*-retinal (Cayman Chemical) at least two days before experiments. All experiments were conducted in the period of 1 hour after lights-on and 1 hour before lights-off (Zeitgeber time 1-11hrs). For all behavioral experiments, mixed populations of male and female flies aged 3-10 days-old were used. For all imaging and functional connectivity experiments, 5-8 day old females were used. A table at the end of this section lists all genotypes used in all experiments.

### Calcium imaging during classical conditioning

Each 5-8 day-old female fly was aspirated without anesthesia into a custom-made chamber comprising a plastic slide, a 200uL pipette tip, and some copper foil tape (Tapes Master). The pipette tip acts as a chamber for the fly, and the copper tape is connected via conductive epoxy (Atom Adhesives) to wires which connect to the power supply to deliver electric shocks. In this setup, the fly’s head was accessible for dissection, its body was contained within the pipette tip, and the back of its thorax was in contact with the copper tape. A thin layer of conductive gel (Parker Labs) was applied to the copper tape prior to inserting the fly, to improve conductivity. The fly was then head-fixed using two-component epoxy glue (Devcon); its proboscis was also glued, but its body and legs were free to move. After the glue dried (about 5 minutes), the cuticle above the head was dissected and air sacs were removed using a 30-gauge syringe needle (Covidien) and a pair of forceps (Fine Science Tools). The head capsule was then sealed with silicone adhesive Kwik-Sil (World Precision Instruments). The fly was then placed in a humidified chamber to recover for 15 minutes.

The fly was then positioned under the microscope. The odor tube was placed about 5mm away from the fly. The odorants used were 0.1% 3-octanol and 0.1% 4-methylcyclohexanol (Sigma Aldrich), controlled via three-way solenoids (Lee Company). The flow rate that was kept constant at 1000mL/min for each odor stream using mass flow controllers (Alicat Scientific). Before reaching the fly, the odor delivery line was split into two, one directed to the fly and the other directed to the photoionization detector (mini-PID, Aurora Scientific) to monitor odor delivery for each trial, for a final flow rate of 500mL/min at each end. The fly was allowed to acclimate to the airflow for at least one minute before beginning the experiment.

Each trial lasted 1 minute; during the first 20s of each trial, blue and green lights were presented simultaneously to visualize fluorescence. The imaging experiment began with four such trials, for the animal to acclimate to the lights. For the actual experiment, alternating trials of 5s of each odor was presented 5s after the light onset. On trials where electric shock was delivered, shock was presented 4s after odor onset for 100ms at 120V.

### Calcium imaging specifications & analysis

Imaging was performed on a Zeiss Axio Examiner upright microscope using a 20x air objective (Zeiss). Using a Colibri LED system (Zeiss), GCaMP was excited at 470nm (blue) and tdTomato was excited at 555nm (green). An optosplitter was used to acquire fluorescence from both wavelengths to allow for ratiometric analysis for fluorescence, to control for movement artefacts. Images were acquired at 5fps.

Data acquired from calcium imaging were analyzed using a combination of Zen (Zeiss) and custom MATLAB code (Mathworks). Since tdTomato was expressed in the same neurons that GCaMP was expressed in, regions of interest (ROIs) were manually drawn based on the tdTomato image and applied to the corresponding GCaMP frame. Mean pixel intensity within each ROI was extracted as raw fluorescence value representing signal in each frame. The GCaMP signal was divided by the tdTomato signal to form the ratiometric fluorescence value, and then processed with a double-exponential fitting to compensate for any photobleaching. The resultant trace was converted into ΔR/R0 by using a baseline intensity (R0), defined as mean intensity during the first 5s of recording.

Mean odor response was defined as the mean ΔR/R0 during the first four seconds of odor onset (*t* = 0-4s); this was so that the response would not be confounded by any potential electric shock response (that would occur 4s after odor onset). Mean shock response was defined as the mean ΔR/R0 during 800ms upon electric shock onset (*t* = 4-4.8s). This again was to avoid confounding this response with any potential odor offset response that would occur at *t* = 5s. In screen data, “difference in mean odor response” refers to the difference (subtraction) of mean odor response on the 5th acquisition trial and the 1st acquisition trial. For reversal trials, this refers to the 2nd reversal trial minus the 1st reversal trial. In data for individual neurons, “relative odor response” is the difference in mean odor response relative to the mean odor response on the first training or reversal trial. Positive values indicate an increase in odor response.

### Behavioral classical conditioning assays

Each group of 30-70 flies was aspirated into a training chamber (Con-Elektronik), comprising an electric shock system and an odor delivery system. The odorants used were 0.07% 3-octanol and 0.1% 4-methylcyclohexanol. The odors were delivered at a flow rate of 500mL/min. Flies were allowed to acclimate in the training chamber for about a minute, and then training began. Odorants were presented for 20s, with 20s inter-trial interval in which clean air was presented. Depending on the protocol, electric shocks may have been delivered during odor presentation. These shocks were 120V for 1.5s, with an inter-trial interval of 3.5s; four of them were delivered during each 20s odor pulse. Within a minute after training, flies were aspirated into the testing arena (Klapoetke et al., 2014) and allowed to walk around in the flat cylindrical arena with CS+ and CS-odors delivered to alternating quadrants at a flow rate of 100mL/min each. Videography was performed under IR light using a camera (Flea3 USB 3.0 camera; Point Grey). CS+ avoidance was quantified as the fraction of flies in the CS-quadrants after being allowed to explore for 2min. Specifically, it was calculated as [(#flies in CS-quadrants) – (#flies in CS+ quadrants)] / (total # flies); an avoidance index of 1 indicates that 100% of the flies chose CS-quadrants, and an avoidance index of 0 indicates that flies chose the two odors equally. Each odorant was alternately used as CS+ and CS-odors in separate groups of flies trained and tested contemporaneously and averaged to generate a reciprocally balanced preference score.

### Optogenetics in conjunction with behavioral classical conditioning assays

The identical setup as described above was used, with the addition of a custom-made RGB LED board positioned ∼3cm below the training chamber. Because the electric shock training chamber was translucent, light was able to pass through and illuminate the chamber. The high-power LEDs (Luxeon) were red (617nm, 166lm) and green (530nm, 125lm), and were controlled together with the odor and electric shock stimuli via a microcontroller (Arduino Mega). Unless otherwise stated, red and green light were set to 50% and 100% max intensity respectively; both were pulsed at 50Hz with a duty cycle of 50%.

### Innate quadrant preference assay

The choice assay was performed in the testing arena described above, but without any odor input. Approximately 30 flies were aspirated into the arena in the dark for one minute before the experiment commenced. For 30s, two diagonal quadrants were illuminated with 627nm red LEDs to activate Chrimson-containing neurons, with no illuminated quadrants 30s after. This was then repeated with the other two quadrants. The fraction of flies (as calculated above) on illuminated quadrants from these two experiments was averaged to calculate the red light preference index. This was then repeated, often with the same group of flies, over a range of light intensities: 1%, 5%, 10%, 20% & 50%.

### Functional connectivity experiments

These experiments were performed similarly to the calcium imaging experiments described in the previous section, with the following modifications: After being placed under the microscope and focusing the field of view, each fly was given 5 minutes in darkness to recover. For direct stimulation experiments (**Figure 4G**,**H**), activity was recorded for 20s without any red light. After 5 minutes, we recorded 10s of baseline activity using dim blue light (470nm at 20% intensity) before stimulating with red light (590nm at 50% intensity) for 500ms, recording for a total of 20s. For experiments looking at odor-evoked responses (**Figure 2H**,**I**), a total of three odor pulses (of 0.01% 4-methylcyclohexanol) were delivered at 2min intervals. Each 5s odor pulse was preceded by 10s of baseline recording (using dim blue light at 20% intensity), and recorded for 5s after odor offset (20s total per trial). During the second odor presentation, a red light pulse (590nm) was delivered for 3s at 8Hz, beginning 0.5s after odor onset. (Blue light was not presented during this trial.) For experiments looking at optogenetic stimulation after aversive memory acquisition (**Figure 5H**,**I**), the protocol was identical to that used in the calcium imaging screen, except only recording (i.e., shining blue light) during the pre-acquisition CS+ trial, the 5^th^ CS+ acquisition trial, and the 2^nd^ CS+ reversal trial. Optogenetic stimulation on the first reversal trial was done using a 590nm red light at 50% intensity, for 3s at 8Hz beginning 1.5s after CS+ onset. Mean odor responses were calculated as mean ΔR/R for the 5s duration that the odor was presented.

### Anatomical synapse analysis

Publicly available, open source electron microscopy data was utilized for all analyses. Details regarding the generation and initial analysis of the primary data can be found in the original report (Scheffer et al., 2020; Xu et al., 2020). The primary dataset utilized in this analysis predominantly imaged one-half of a female *D*rosophila brain, as such we limited our analysis to this side of the data and included only neurons that were marked as fully traced to avoid biasing in our sampling. We first manually identified neurons in the neuPrint database that corresponded to our neurons of interest. Neuronal skeletons were registered to the JRC2018F template brain and visualized in three-dimensions using natverse R packages (Bates et al., 2019). After identification, the number and ROI of synaptic contacts between neurons of interest were quantified using existing neuPrint Python API functions for neuPrint. Specific three-dimensional Euclidean coordinates of postsynaptic contacts of interest were generated and visualized on particular postsynaptic neurons in two-dimensions using existing neuPrint functions.

### Statistical analyses

Statistical analyses were performed in GraphPad Prism 8. All data was tested for normality using the D’Agostino and Pearson omnibus test. Each statistical test used in each figure is reported in the legends. Generally, normally distributed data were analyzed with a *t*-test or one-way ANOVA followed by Dunnett’s post-hoc test where appropriate. For non-Gaussian distributed data, a Wilcoxon signed-rank test, Mann-Whitney test, or Kruskal-Wallis test was performed followed by Dunn’s multiple comparisons test where appropriate.

**Table 1:**
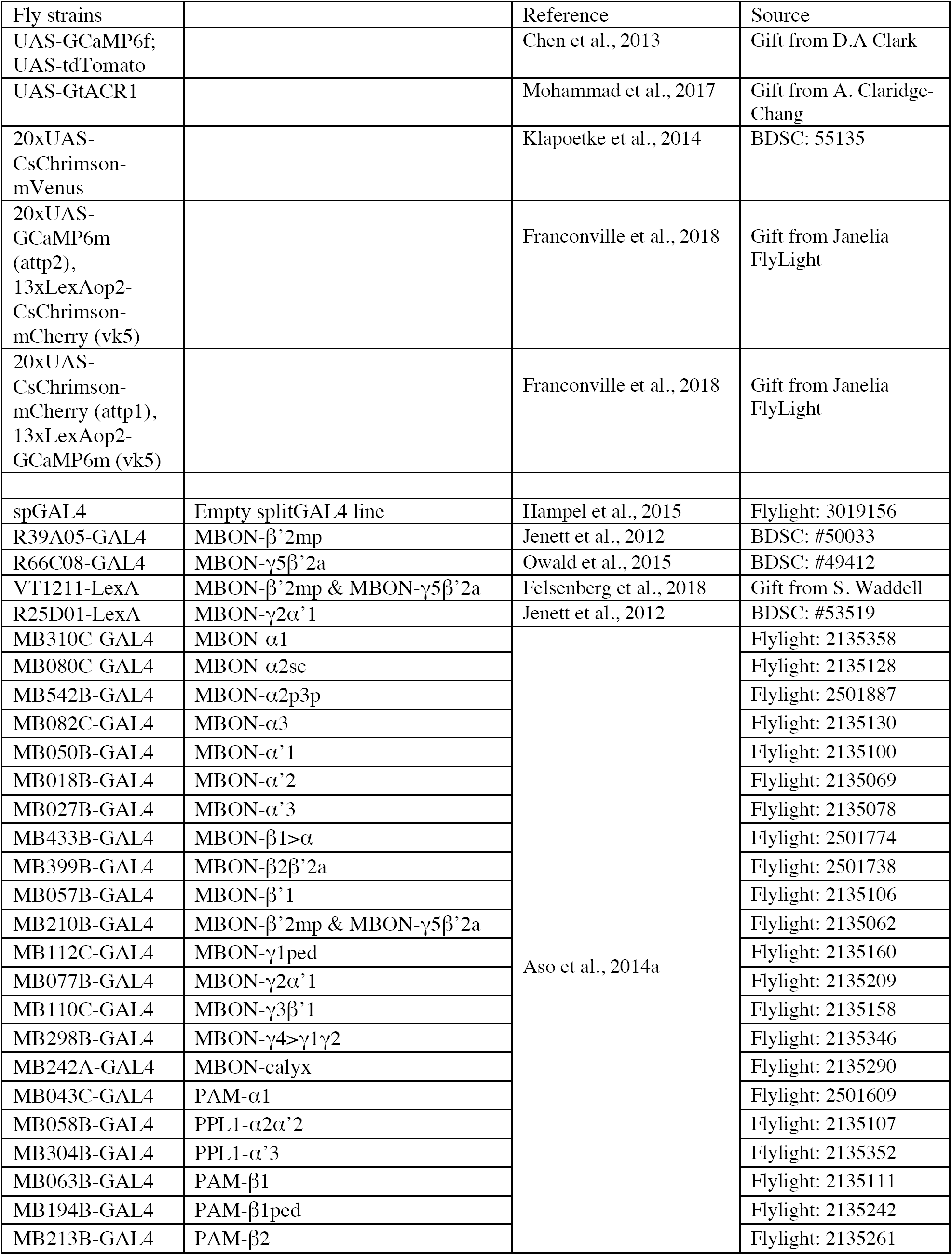
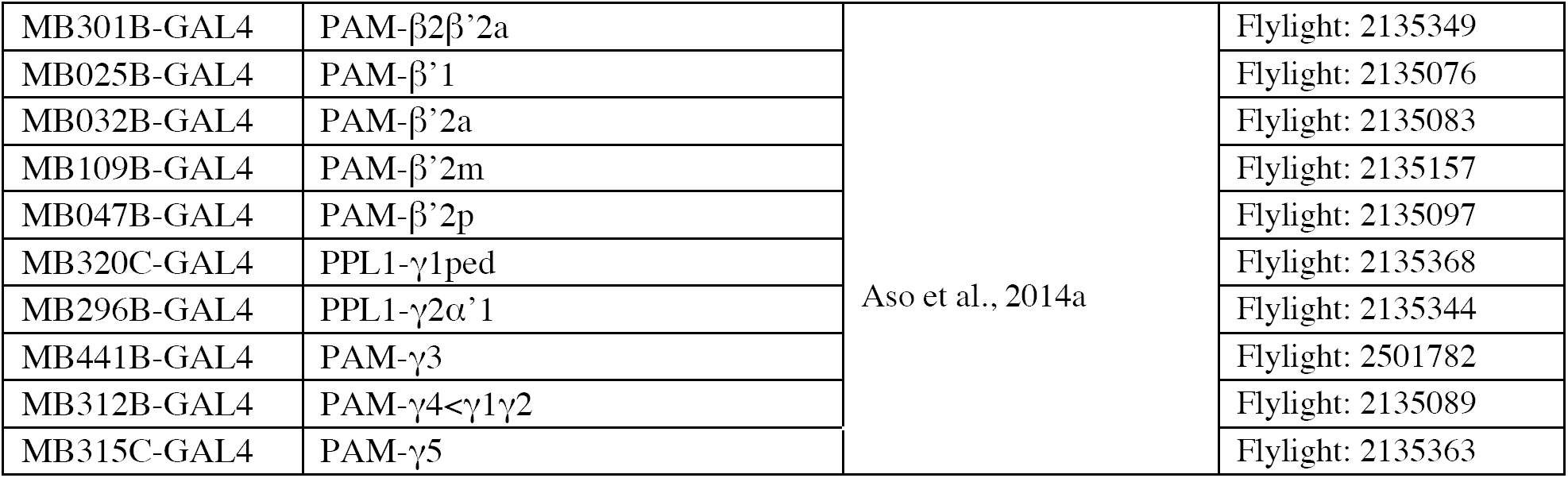
List of fly strains.

## Author Contributions

Designed study and experiments: L.Y.M, M.N.N.

Contributed unpublished reagents: P.S.

Performed experiments: L.Y.M.

Data analysis and statistics: L.Y.M, P.D., M.N.N.

Wrote the paper: L.Y.M., M.N.N.

## Acknowledgments

We thank Damon Clark and Scott Waddell for reagents. We thank members of the Nitabach lab for technical support, advice and comments. We thank Zunaira Arshad, Alex Buhimschi, Juliana Rioz Chen, Justin Choi, Michelle Benavidez Frausto, Maddie Gharizan, Daniel Grubman, Kaitlyn Kang, Dawit Mengasha, Alice Oh, Natalie Orner, and Krishna Vali for conducting pilot experiments.

L.Y.M. was supported by the National Science Foundation (NSF) Graduate Research Fellowship Program (GRFP), Yale University Cellular and Molecular Biology training grant from the National Institutes of Health (NIH) (T32GM007223), Gruber Science Fellowship, and National University of Singapore Overseas Graduate Scholarship. P.A.D is supported by the Medical Scientist Training Grant, NIH (T32GM007205). Work in the laboratory of M.N.N. is supported by National Institute of Neurological Disease and Stroke (NINDS), NIH (R01NS091070) and National Institute of General Medical Studies (NIGMS), NIH (R01GM098931).

---

**Supplemental Figure 1:**
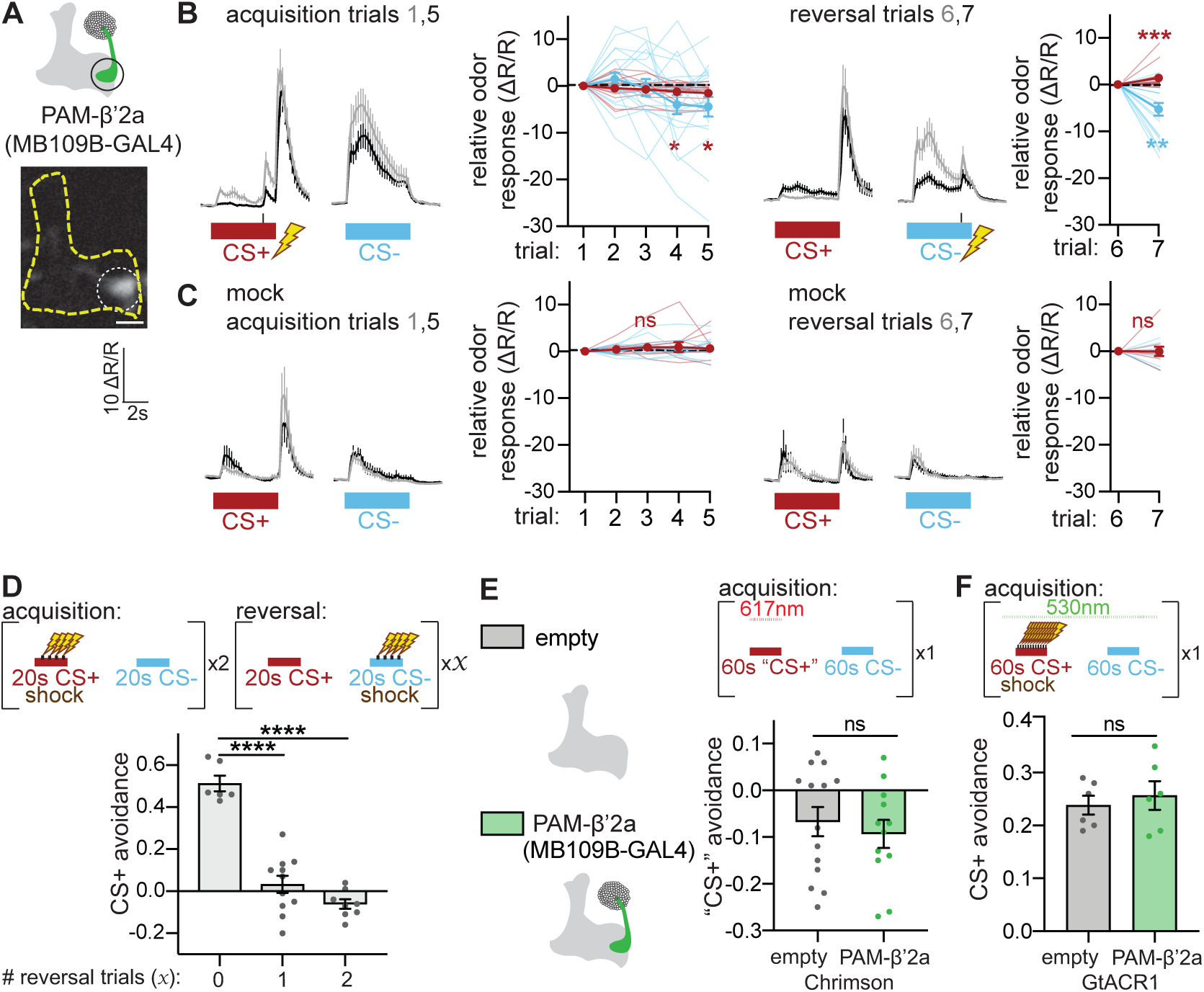
Related to Figure 1. (**A**) Schematic and sample fluorescence image of PAM-β’2a. (**B**) Neural activity of PAM-β’2a in flies undergoing aversive memory acquisition and reversal trials, using reciprocal odor identities, i.e., 3-octanol and 4-methyl-cyclohexanol as CS+ and CS-odors respectively. PAM-β’2a CS+ odor response decreases during acquisition and increases during reversal. *n* = 19 flies. Statistical comparisons as in **Figure 1F**. (**C**) Same as in (**B**), but from flies undergoing mock aversive memory acquisition and reversal, i.e., odors are presented as in (**B**), but no shocks are delivered. No change in PAM-β’2a odor response occurs during acquisition or reversal. *n* = 11 flies. Same statistical comparisons are used as in (**B**). (**D**) Top: Reversal protocol for behavioral experiments. During acquisition, 20s of CS+ odor is paired with four electric shocks, followed by 20s of CS-odor without reinforcement; this is done twice. After acquisition, flies undergo either zero, one or two reversal trials, in which 20s of CS+ odor is presented without electric shock, and 20s of CS-odor is presented with electric shock. Flies are then placed in quadrant arena and odor preference (CS+ vs CS-) is determined. Bottom: Flies that do not undergo reversal after acquisition robustly avoid CS+ odor. Flies that undergo one or two reversal trials exhibit greatly reduced CS+ avoidance. *n* = 6, 11 & 8 groups of flies per condition respectively. Statistical comparison is by one-way ANOVA with Dunnett’s post-hoc test for multiple comparisons. (**E**) Top: Experimental protocol for assessing valence associated with odor when paired with red-light neural activation via Chrimson. Flies are placed in a shock tube and exposed to 1min pulse of “CS+” odor paired with red light illumination of the shock tube to activate neurons expressing Chrimson, followed by a 1min pulse of “CS-” odor without red light. Flies are then placed in quadrant arena and odor preference was determined. Bottom: Flies expressing Chrimson in PAM-β’2a do not increase preference for “CS+” odor, relative to control flies. *n* = 14 & 12 independent groups of flies per genotype. Statistical comparison is by unpaired *t*-test. (**F**) Above: Experimental protocol for assessing necessity of neurons during aversive memory acquisition using green-light neural silencing via GtACR1. Flies undergo aversive memory acquisition protocol in which 1min of CS+ is paired with 12 electric shocks; 1min of CS-is presented without reinforcement. The shock tube is illuminated with green light throughout acquisition. Below: Control flies acquire a robust aversive memory associated with the CS+ odor and avoid CS+ quadrants. Aversive memory acquisition is not impaired in flies expressing GtACR1 in PAM-β’2a. *n* = 6 independent groups of flies per genotype.

**Supplemental Figure 2:**
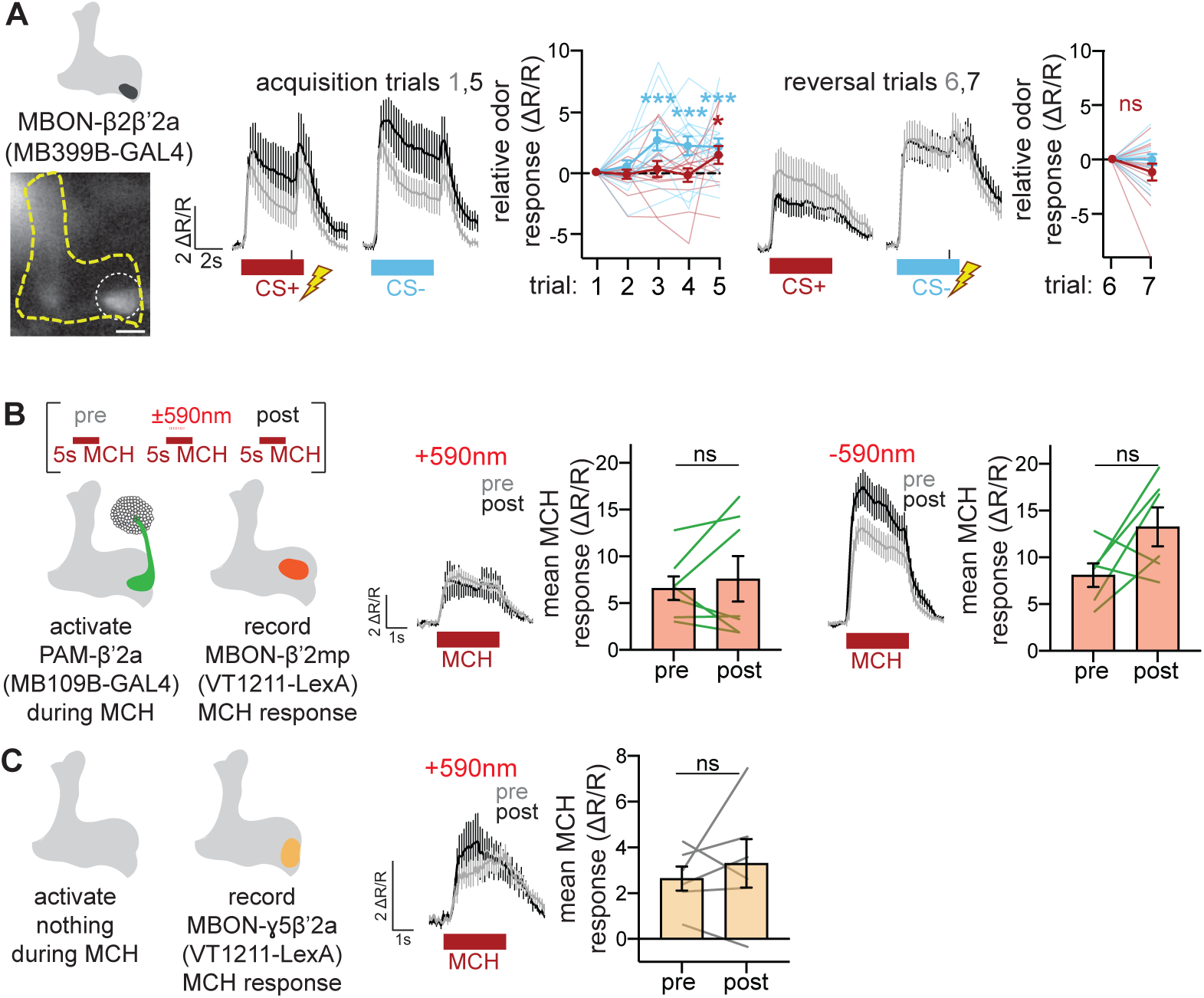
Related to Figure 2. (**A**) Neural activity of MBON-β2β’2a in flies undergoing aversive memory acquisition and reversal trials. Both CS+ and CS-odor responses increase in MBON-β2β’2a during acquisition. Importantly, no changes in odor response occur during reversal, making it an unlikely candidate as the postsynaptic partner of PAM-β’2a that decreases CS+ avoidance behavior. *n* = 14 flies. (**B**) Same experiment as in **Figure 2G**. Neural responses in MBON-β’2mp before (grey) and after (black) odor-light pairing. Optogenetic activation of PAM-β’2a during MCH odor delivery does not affect subsequent MBON-β’2mp MCH odor response. This is also true in the mock condition. *n* = 7 & 6 flies. Statistical comparison is by paired *t*-test. (**C**) Same experiment as above, except without expressing Chrimson in any neurons. Odor responses in MBON-γ5β’2a before (grey) and after (black) odor-light pairing are not significantly different. *n* = 6 flies. Statistical comparison is by paired *t*-test.

**Supplemental Figure 3:**
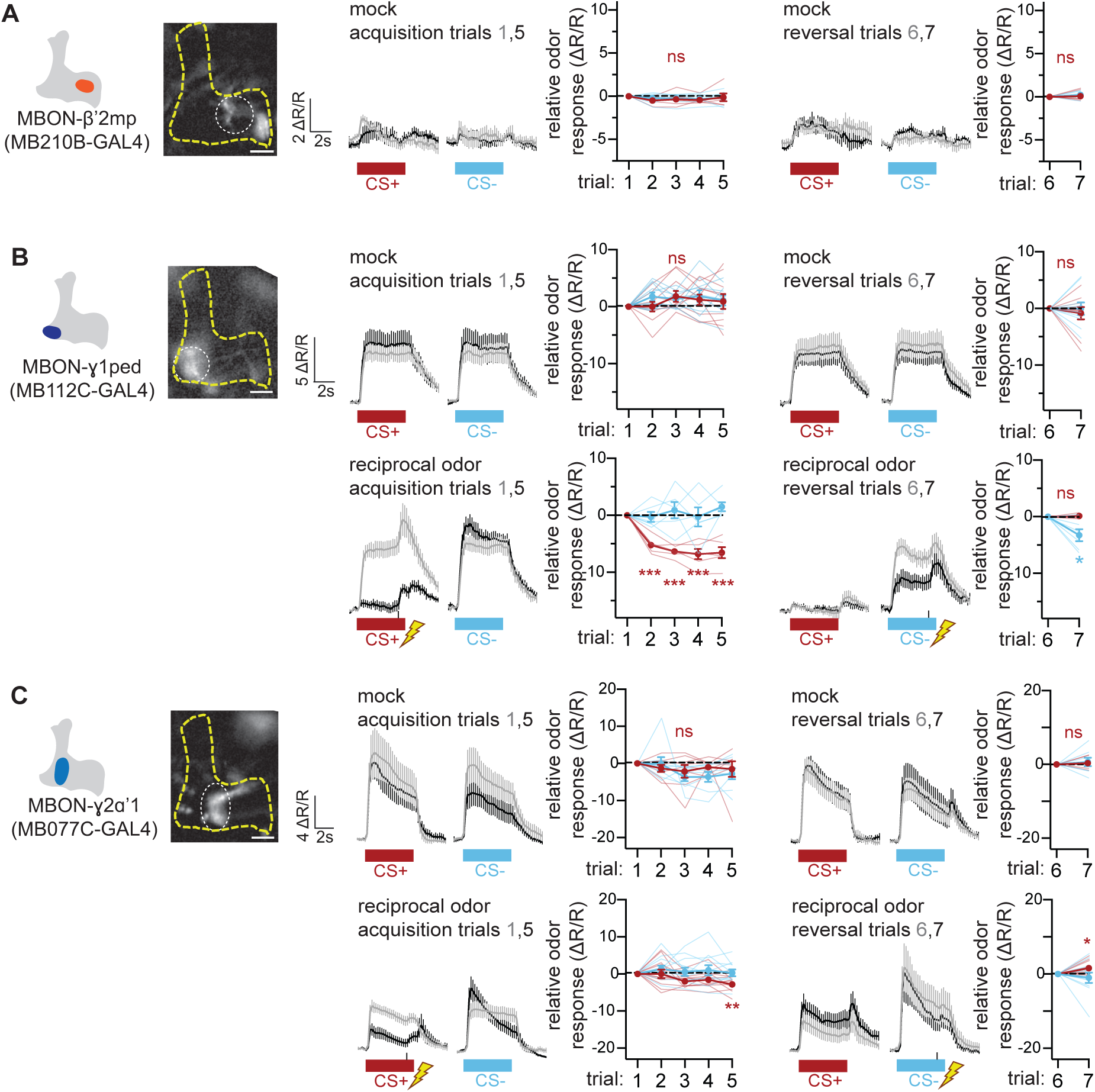
Related to Figure 3. (**A**) No change in MBON-β’2mp odor response occurs during mock acquisition or reversal. *n* = 6 flies. (**B**) No change in MBON-γ1ped odor response occurs during mock acquisition or reversal. CS+ odor response decreases during acquisition, and CS-odor response decreases during reversal using reciprocal odors (i.e., 3-octanol and 4-methylcyclohexanol as CS+ and CS-respectively). *N* = 10 & 6 flies. (**C**) No change in MBON-γ2α’1 odor response occurs during mock acquisition or reversal. CS+ odor response decreases during acquisition, and increases during reversal using reciprocal odors. *n* = 8 & 11 flies.

**Supplemental Figure 4:**
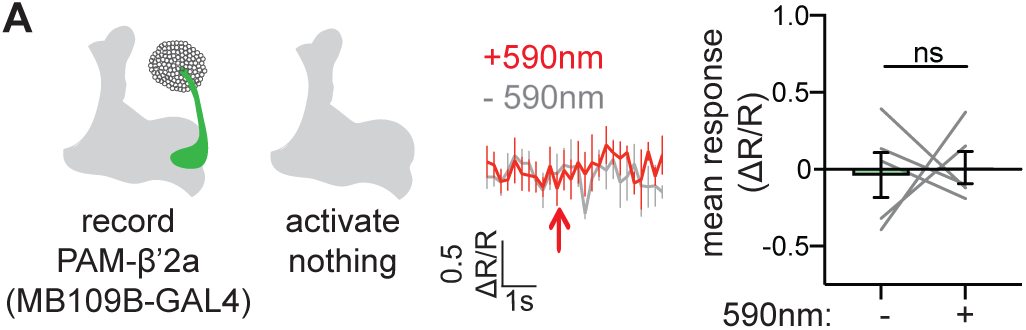
Related to Figure 4. (**A**) Same experiment as in **Figure 4G**, except without expressing Chrimson in any neuron. PAM-β’2a activity does not change during red light illumination. *n* = 5.

**Supplemental Figure 5:**
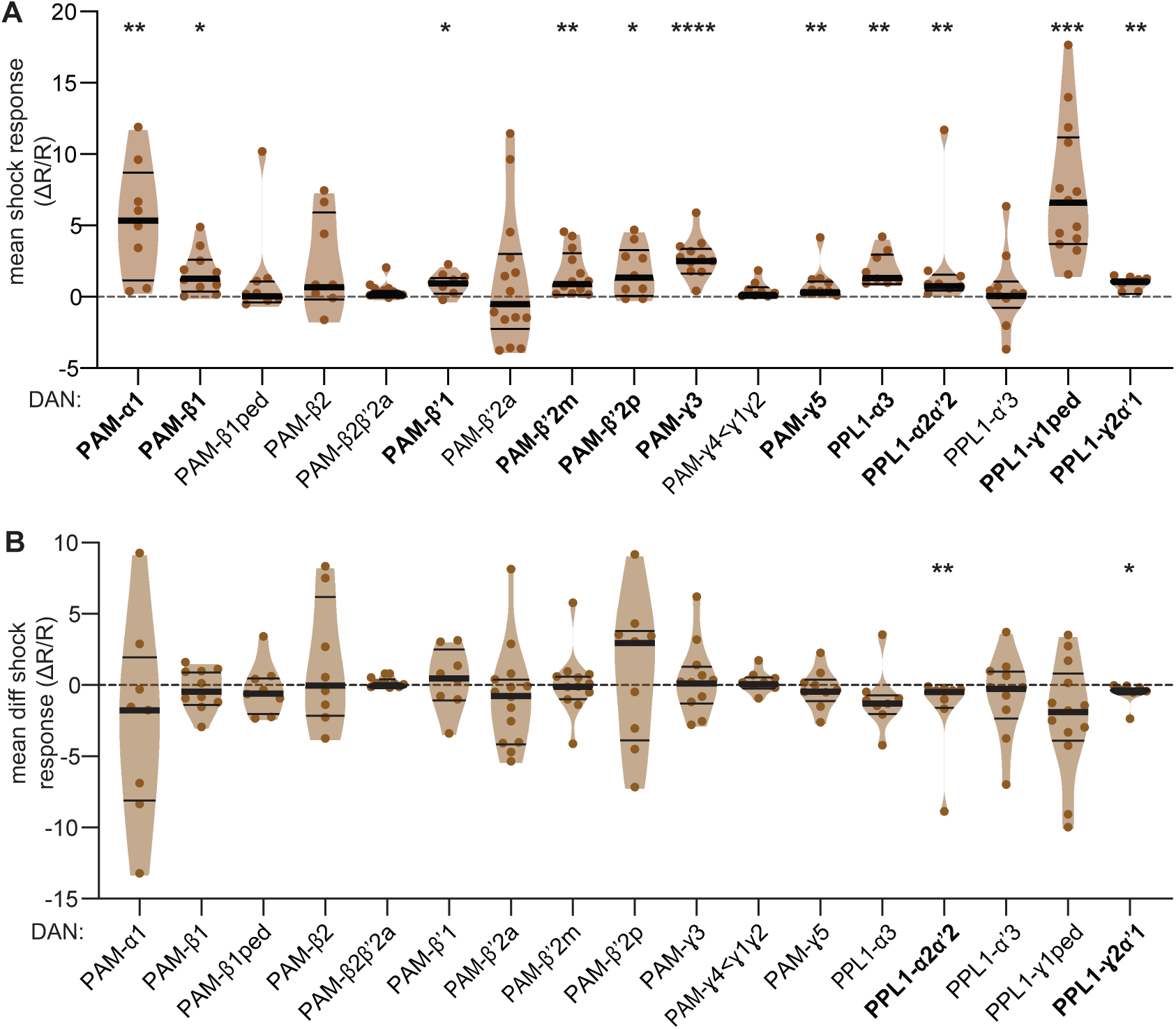
Related to Figure 5A. (**A**) Mean shock response of each DAN subset during aversive memory acquisition portion of imaging screen. Mean shock response is defined as average activity starting 800ms from electric shock onset, averaged across all 5 acquisition trials. Since CS+ odor offset occurs 1s after shock onset, the mean shock response is defined as such to avoid the time window during odor offset to prevent confounds. Violin plots show range of data points, with median (thick black line) and quartiles (thin black lines). Most PPL1 DANs respond to electric shock, as expected. Surprisingly, many PAM DANs also respond to electric shock. *n* = 7-12 flies per genotype. Statistical comparison is by uncorrected one-sample *t*-test or Wilcoxon signed-rank test for each genotype. (**B**) Difference in mean shock response between first and last acquisition trials. Positive values indicate a relative increase in shock response over the course of acquisition. *n* = 7-12 flies per genotype. Statistical comparison is by uncorrected one-sample *t*-test or Wilcoxon signed-rank test for each genotype.

**Supplemental Figure 6:**
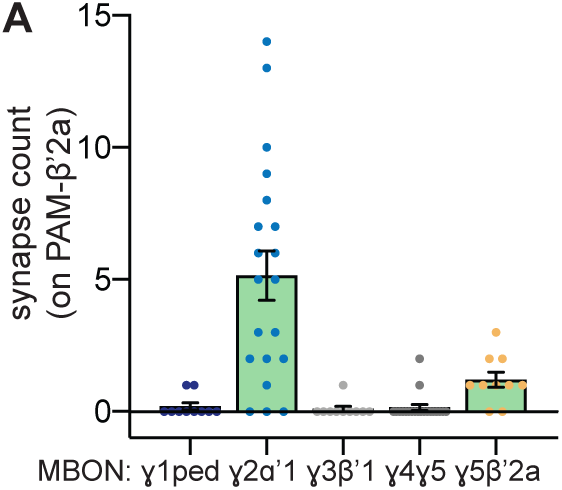
Related to Figure 5G-I. (**A**) Number of synapses between each γ lobe MBON and PAM-β’2a. MBON-γ2α’1 synapses plotted again (as in **Figure 5H**) for ease of comparison. MBON-γ2α’1 has more synaptic connections on PAM-β’2a than any other γ lobe MBON.

